# Engineering a Biomimetic Multiphasic Suture Anchor System for Enhanced Rotator Cuff Enthesis Regeneration

**DOI:** 10.1101/2025.10.30.685549

**Authors:** Zizhao Li, Se-Hwan Lee, Lin Xu, Marina Santos, Ruqiang Lu, Ellen Y. Zhang, DongHwa Kim, Bat-Ider Tumenbayar, Tyler Blanch, Jaeun Jung, Jayden Shin, Yongho Bae, Richard T. Tran, Thomas Schaer, Su Chin Heo

**Affiliations:** McKay Orthopaedic Research Laboratory, Department of Orthopaedic Surgery University of Pennsylvania, 3450 Hamilton Walk, Philadelphia, PA, 19104, United States; Department of Bioengineering, University of Pennsylvania, 210 South 33rd Street, Philadelphia, PA, 19104, United States; New Bolton Center, Department of Clinical Studies, University of Pennsylvania, 382 West Street Road Kennett Square, Philadelphia, PA, 19348, United States; Acuitive Technologies, Inc., 50 Commerce Drive, Allendale, NJ, 07401, United States; Department of Pathology and Anatomical Sciences and Department of Biomedical Engineering, University at Buffalo, 955 Main Street, Buffalo, NY, 14203, United States

**Author notes:** **Funding:** Foundation for the National Institute of Health (K01 AR07787, R21 R077700, P30 AR069619, R01 HL163168), National Science Foundation (CMMI 1548571), 2023-2024 University of Pennsylvania Health-Tech Accelerator Award.

**Keywords:** Rotator Cuff Repair, Enthesis Regeneration, Tunable decellularized extracellular matrix, citrate-based polymer, Multiphasic scaffold

## Abstract

Conventional suture anchor methods in rotator cuff repair often fail to replicate the native tendon-to-bone interface, leading to re-tears due to stress concentrations and poor biological integration at anchor sites. To address these challenges, we engineered a biomimetic multiphasic scaffold system (BMS) that integrates with standard suture anchors and deliver spatially organized structural and biological cues to enhance enthesis regeneration. The BMS comprises three distinct phases: aligned nanofibrous decellularized bovine Achilles tendon extracellular matrix (dECM) with ‘stiff’ methacrylated hyaluronic acid (MeHA) for tendon regeneration; nonaligned nanofibrous dECM with ‘soft’ MeHA for fibrocartilage formation; and a porous, citrate-based composite scaffold with bioactive glass for bone integration. *In vitro*, the BMS facilitated zone-specific tenogenic, fibrochondrogenic, and chondrogenic differentiation. Further, *in vivo*, it promoted successful integrative healing, forming distinct tendon, fibrocartilage, and bone regions at the repair site. This advanced multiphasic scaffold replicates native tissue properties, offering a promising strategy to improve rotator cuff repair. Its integration with conventional suture anchors provides an innovative design that enhances mechanical fixation and guides enthesis healing to reduce re-tear rates. Broadly, this platform offers a versatile solution for biointegrative repair strategies across complex soft-to-hard tissue interfaces.

## 1. Introduction

The rotator cuff comprises four muscles and their associated tendons which stabilize the joint and enable shoulder movement ^[1,2,3]^. Rotator cuff tears lead to significant pain, weakness, irritation, and diminished range of motion in affected individuals (**Figure 1a**) ^[1,2,3]^. These injuries are particularly prevalent in the adult population, with approximately 2 million cases reported annually in the United States and 30% of adults over 60 years old, and 62% of adults over 80 years old, affected globally ^[4,5]^. Current arthroscopic shoulder repairs use suture anchor systems to reattach the torn tendon to the humeral head ^[6,7]^. However, these procedures often fail to restore the native enthesis structure, resulting in scar tissue formation, reduced mechanical integrity, and a high incidence of re-tear (**Figure 1a**) ^[8,9]^. A key limitation lies in the design of conventional suture anchors, typically made of bioinert polymers which lack the biochemical and mechanical cues necessary to support enthesis regeneration ^[10,11]^. These challenges emphasize an urgent need for advanced therapeutic strategies that improve functional and durable tendon-to-bone healing.

**Figure 1.**
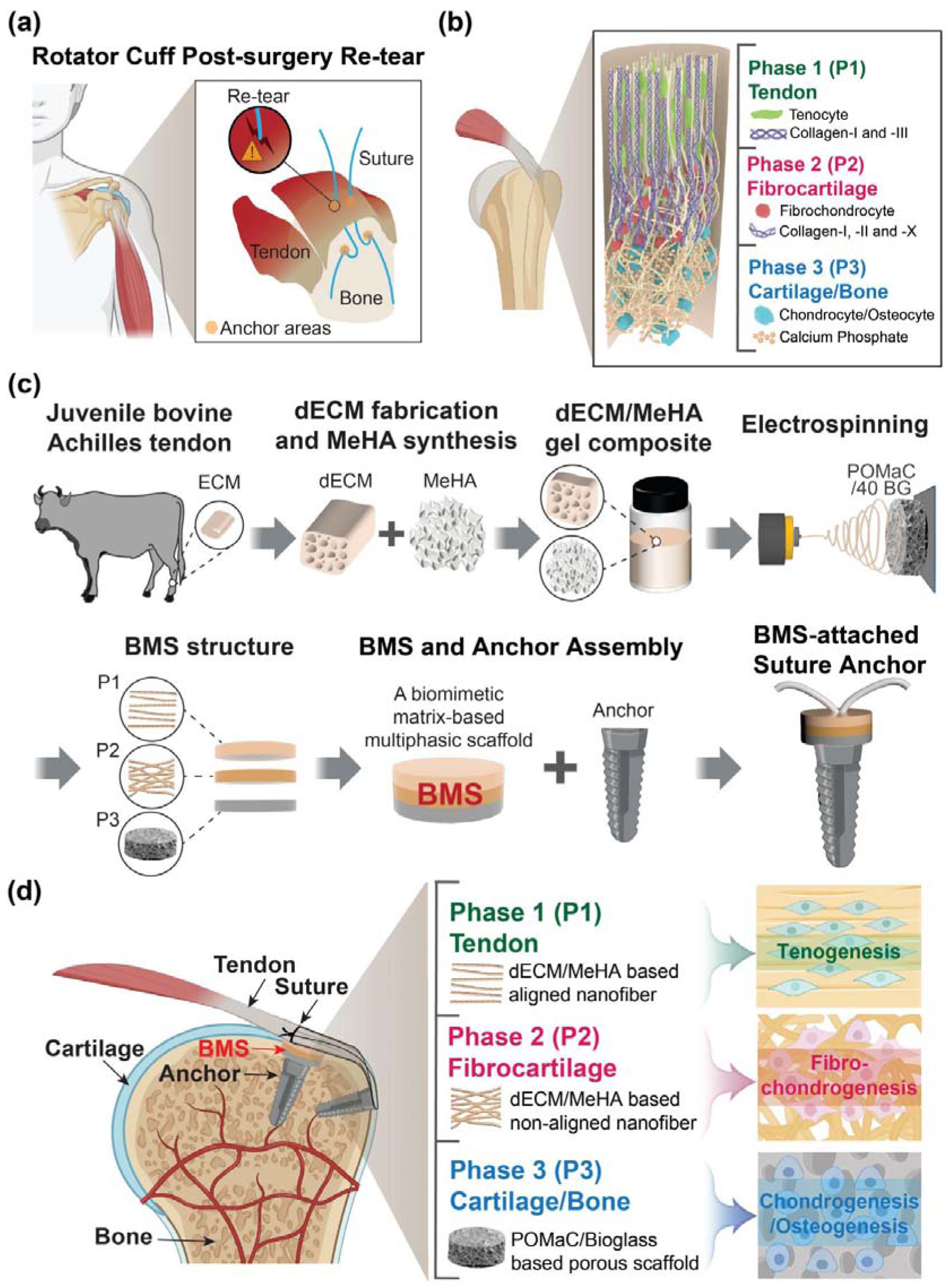
Engineering a biomimetic, multiphasic suture anchor system for enhanced rotator cuff repair. (a) Schematic of rotator cuff repair, emphasizing the potential for re-tear at suture sites post-surgery. (b) Structural organization of the native rotator cuff tendon-to-bone insertion (enthesis) provides insight into the suture anchor-attached biomimetic multiphasic scaffold (BMS) design. (c) Schematic diagram illustrating the strategy for engineering the BMS to be used with suture anchors. (d) Schematic illustration of the BMS scaffold integrated with suture anchors in rotator cuff repair techniques to enhance enthesis regeneration.

The native rotator cuff enthesis is a complex, multi-tissue interface characterized by a seamless transition from tendon to fibrocartilage to bone (**Figure 1b)** ^[4,5]^. The zone-dependent architecture of the enthesis is crucial for reducing stress concentrations, enabling effective load transfer, and providing mechanical stability, yet remains difficult to regenerate. While recent regenerative engineering advances have led to the development of commercial scaffolds ^[12,13,14,15]^, challenges persist. Notable commercial examples include porous implants derived from bovine Achilles tendon and polylactide fiber-based scaffolds designed to support healing ^[16,17]^. However, these scaffolds often do not replicate the multi-tissue architecture and lack key biochemical cues, limiting their effectiveness in restoring the native enthesis. Thus, a biomimetic approach that recapitulates these critical features is essential for improving rotator cuff repair outcomes.

Decellularized extracellular matrix (dECM) is a versatile and promising native biomaterial for tissue-specific applications ^[18]^. Its natural composition provides structural integrity and essential tissue-specific biochemical cues, making it ideal for regenerative engineering strategies ^[19,20,21]^. Building on our previous work, which demonstrated the efficacy of dECM-based nanofibrous scaffolds for fibrous connective tissue regeneration ^[22]^, in this study, we have developed a significantly advanced biomimetic, dECM-based multiphasic scaffold system (BMS). This novel system integrates bovine Achilles tendon-derived dECM with methacrylated hyaluronic acid (MeHA) to promote tendon regeneration ^[23,24]^. The strategic incorporation of MeHA allows for precise control over the hydrogel stiffness ^[23,25]^ to support tendon and fibrocartilaginous regeneration. For fibrocartilage and bone integration, a bioenergetic and biodegradable citrate-based elastomer ^[26,27,28]^, which has been reported to induce osteogenic differentiation in human mesenchymal stem cells ^[26,29]^, is composited with bioactive glass (BG), known for its osteogenic potential ^[30]^. The citrate-based composite provides metabolic and osteogenic cues for fibrocartilage-to-bone integration within the scaffold’s multiphasic structure ^[31,32,33,34]^ (**Figures 1b, c**). Thus, the combination of tendon-derived dECM and MeHA for tendonous repair and a citrate-based elastomeric material with BG for osseous repair recapitulates the zonal differences of the enthesis tissue within our BMS. This multiphasic scaffold seamlessly integrates with suture anchors to improve upon the limitations of existing technologies by replicating the native structural and mechanical properties of the enthesis, thereby enabling phased tissue regeneration (**Figures 1b, c**).

Our BMS consists of three distinct phases, each tailored to the zonal architecture of the enthesis: (1) Phase 1 (P1): aligned, electrospun dECM scaffolds with ‘stiff’ MeHA for tendon regeneration, (2) Phase 2 (P2): nonaligned, electrospun dECM scaffolds with ‘soft’ MeHA for fibrocartilaginous regeneration, and (3) Phase 3 (P3): Poly (octamethylene maleate (anhydride) citrate) (POMaC)/BG porous scaffolds for fibrocartilage-to-bone integration (**Figure 1c**). Comprehensive *in vitro* evaluations demonstrated the zonal functionality of the BMS, with each phase supporting cell morphology, differentiation, and ECM deposition consistent with its intended regenerative role. For instance, ‘stiff’, aligned tendon dECM nanofibrous scaffolds promoted tenogenic differentiation, while ‘soft’, nonaligned dECM scaffolds facilitated fibrochondrogenic differentiation. Additionally, POMaC/BG porous scaffolds demonstrated enhanced mineralization and chondrogenic differentiation, indicating their role in supporting the fibrocartilage-to-bone tissue transition. Further, to evaluate the translational potential of the BMS, we used a rabbit rotator cuff repair model, where native enthesis tissue was surgically removed and replaced with the BMS scaffold. *In vivo* results demonstrated that the BMS facilitated integrative healing, with distinct tendon, fibrocartilage, and bone zones forming at the repair site. Micro-CT analyses revealed enhanced bone remodeling and histology confirmed the successful integration of the scaffold with native tissues, highlighting its potential for zone-specific rotator cuff enthesis repair.

Our multiphasic scaffold system integrates with conventional suture anchors to enhance mechanical fixation and zone-specific regeneration by providing critical biological and structural cues (**Figure 1d**). This biomimetic approach mimics the native enthesis architecture, offering a more effective repair strategy and a strong foundation for future clinical studies.

## 2. Results and Discussion

### 2.1. Decellularized bovine ECM characterization and proteomic analysis

Following the decellularization process, H&E staining confirmed successful decellularization of the bovine Achilles tendon, showing complete removal of nuclei (**Figures S1a, b, Supporting Information**). However, key ECM proteins, specifically collagen and glycosaminoglycans, were retained, as verified by Picrosirius Red and Alcian Blue staining, respectively (**Figure S1b, Supporting Information**). To assess ECM composition before and after decellularization, we conducted proteomic analysis on native and decellularized ECM (dECM) samples. The principal component analysis (PCA) demonstrated distinct clustering of native and dECM samples (**Figure S2a, Supporting Information**). Proteomic analysis identified 2,416 unique proteins in the native bovine Achilles tendon, with 250 proteins common to native and dECM tissues (**Figure S2b, Supporting Information**). Stacked bar plots across four donors (**Figure 2a)** showed overall retention of ECM proteins post-decellularization, with tendon-associated collagens (e.g., COL1A1, COL1A2, COL3A1, COL5A1, COL5A2, COL6A1, COL6A2, COL6A3) preserved in the dECM (**Figure 2a**). We further identified core matrisome components in the dECM and constructed a dense protein-protein interaction network using the STRING database, comprising 52 nodes with 618 protein interactions (**Figures 2b, c**). Remarkably, the dECM retained key ECM proteins, including collagen subtypes (e.g., COL1A1, COL1A2, COL3A1, COL5A1, COL6A1), glycoproteins involved in matrix binding and structural integrity (e.g., LAMC1, LTBP1, ELN, FN1), proteoglycans essential for ECM remodeling and cell-matrix interactions (e.g., BGN, DCN, OGN), and proteins associated with ECM organization and angiogenesis (e.g., POSTN, TNC, THBS4, HSPG2), indicating preserved remodeling capacity, enhanced cell adhesion, and regenerative potential (**Figures 2b, c**). Gene ontology (GO) analysis further revealed a significant enrichment of biological processes related to extracellular structure organization, collagen biosynthesis, and cell-matrix adhesion, with comparable distributions across cellular components and molecular functions in native and decellularized tissues (**Figures 2d and Figures S2c, d, Supporting Information**). These findings indicate that the decellularization process preserved essential ECM components and biological functions of the native tissue, supporting its suitability for regenerative applications.

**Figure 2.**
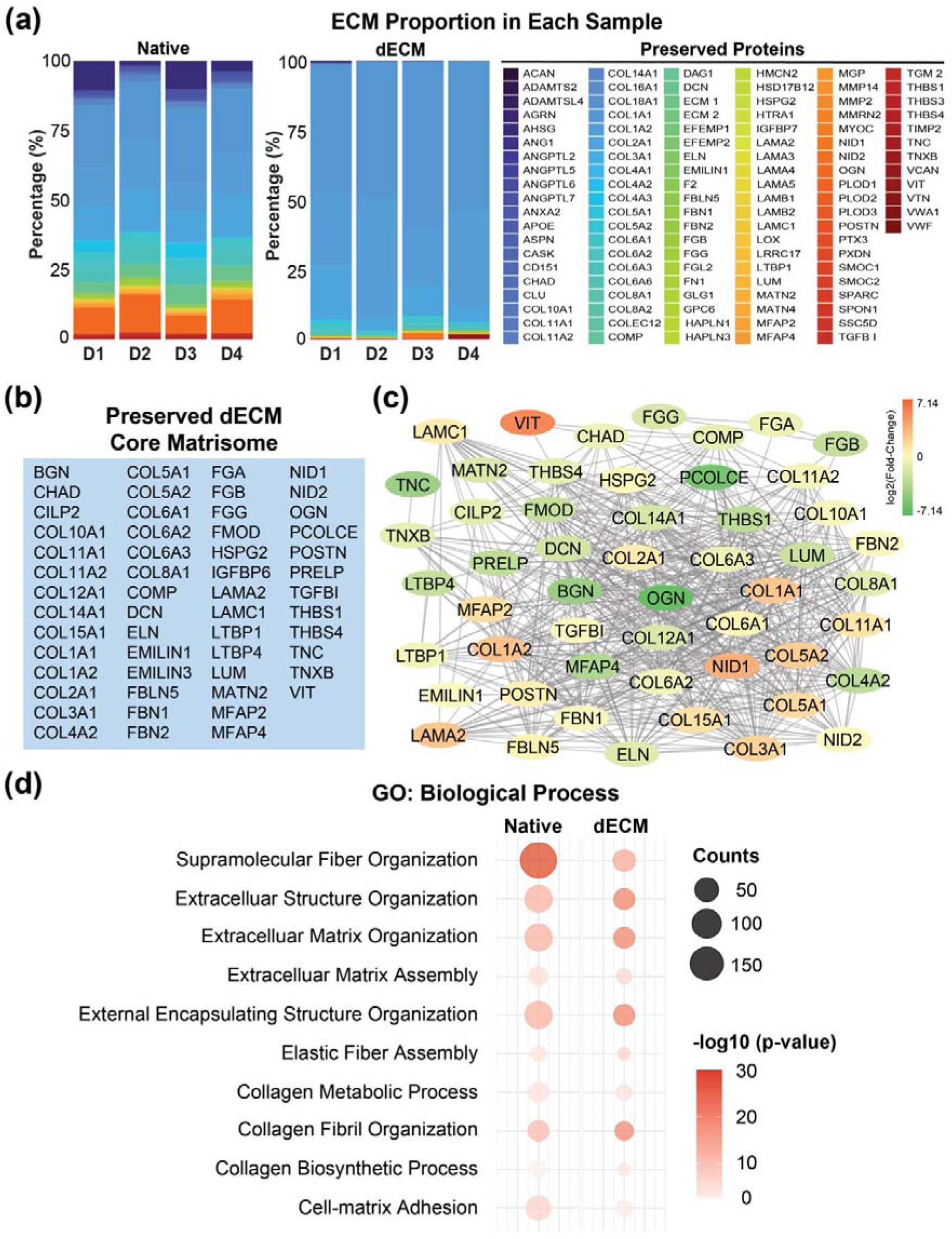
Proteomic profiling of decellularized bovine Achilles tendon ECM. (a) Stacked bar plots showing the composition of ECM proteins in native versus dECM samples across four donors (D = Donor, Donor # = 4/group), highlighting preserved proteins post-decellularization. (b) List of core ECM proteins retained in dECM samples. (c) Protein-protein interaction network illustrating the interconnectivity of core ECM components in dECM. (d) Gene Ontology (GO) analysis of Biological Process categories related to ECM structure and function.

### 2.2. Development and biomechanical characterization of tunable dECM/MeHA nanofiber scaffolds

To develop a tunable tendon dECM-based nanofibrous scaffold with adjustable fiber stiffness and alignment, methacrylated hyaluronic acid (MeHA) was synthesized with a 35% (‘soft’) and 100% (‘stiff’) modification rate ^[23]^ **(Figure S1a, Supporting Information)**. These MeHA variants were blended with dECM, and then electrospun and crosslinked, to create aligned (AL) and non-aligned (NAL) tendon dECM-based nanofibrous scaffolds (**Figure 1c, Figure S1a, Supporting Information**). Scanning electron microscopy (SEM) confirmed the tunable alignment of these nanofibers (**Figure 3a**). Distribution and retention of dECM within the nanofibers was verified via Picrosirius Red, 5-(4,6-dichlorotriazinyl) aminofluorescein (DTAF), and Alcian Blue staining in both ‘stiff’ and ‘soft’ groups, compared to the control (Ctrl) group composed of 100% modified MeHA without dECM (**Figures 3b, c, d**).

**Figure 3.**
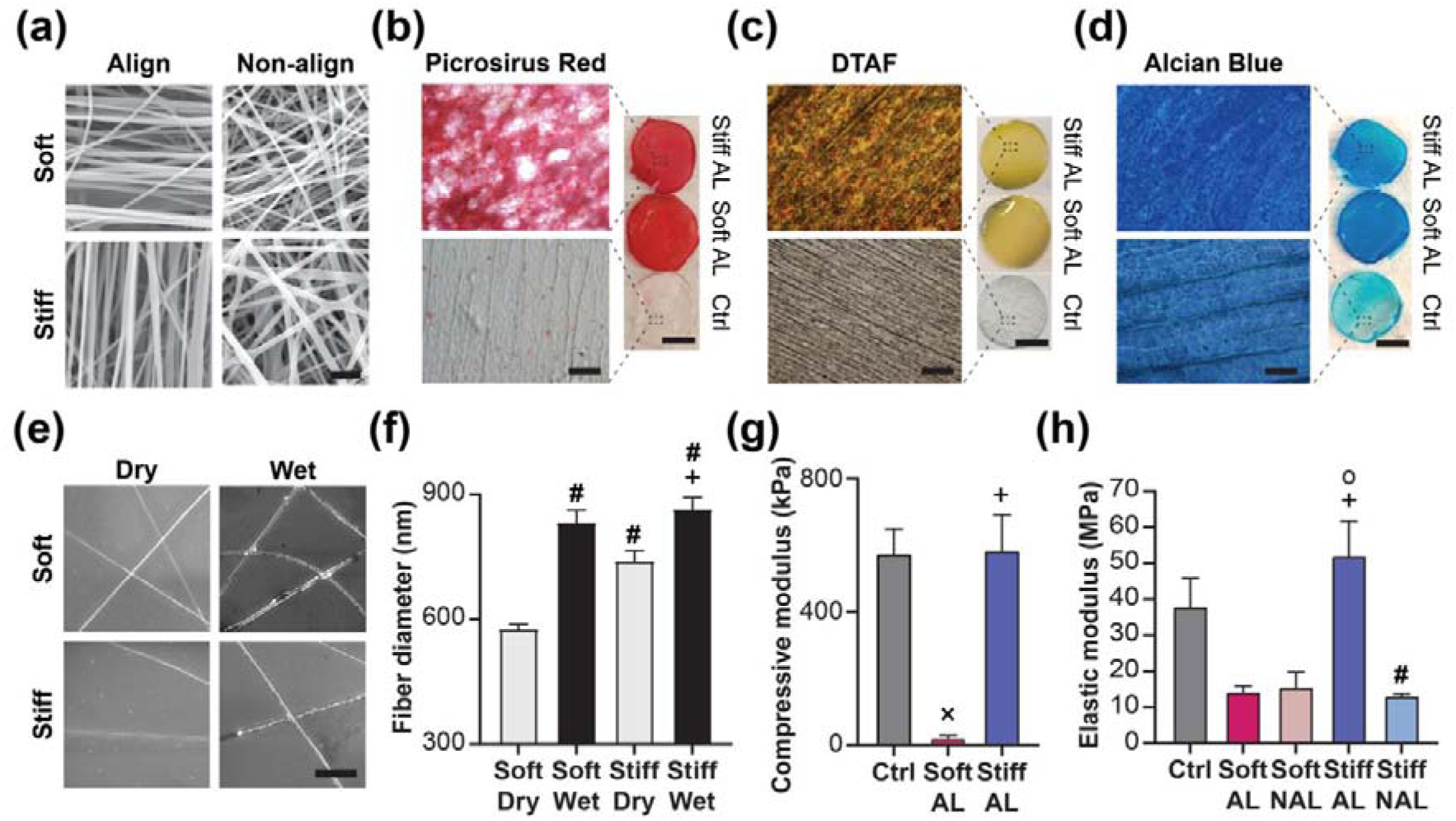
Structural and mechanical characterization of tunable tendon dECM/MeHA nanofiber scaffolds. (a) SEM images of ‘soft’ and ‘stiff’ dECM/MeHA nanofibers (Scale bar = 2 μm). (b) Picrosirius red staining, (c) DTAF staining, and (d) Alcian blue staining of ‘soft’ aligned dECM/MeHA (Soft AL), ‘stiff’ aligned dECM/MeHA (Stiff AL), and ‘stiff’ MeHA-only control (Ctrl) nanofibers (scale bar = 100 μm for enlarged views; 1 cm for circular scaffolds). (e) Representative environmental SEM (E-SEM) images of nanofibers in dry and PBS-swollen (wet) states (Scale bar = 20 μm). (f) Quantification of fiber diameters in dry and wet states (Soft Dry, Soft Wet, Stiff Dry, Stiff Wet; n = 50, #: p < 0.05 vs. Soft Dry; +: p < 0.05 vs. Stiff Dry). (g) Compressive modulus of Ctrl, Soft AL, and Stiff AL nanofibers (n >30, ×: p < 0.05 vs. Ctrl; +: p < 0.05 vs. Soft AL). (h) Elastic modulus of Ctrl, Soft AL, Soft NAL, Stiff AL, and Stiff NAL nanofibers (n =4, ○: p < 0.05 vs. Soft AL; +: p < 0.05 vs. Soft NAL; #: p < 0.05 vs. Stiff AL).

Fiber diameters were measured in dry and PBS-swollen states using environmental SEM (E-SEM, **Figure 3e**). The dry diameter of the ‘stiff’ fibers was approximately 28% larger than the ‘soft’ fibers; however, both ‘stiff’ and ‘soft’ fibers showed a consistent diameter (∼850 nm) when hydrated. The morphological consistency of nanofibers ensures comparable cell-fiber interactions across groups post-hydration, while stiffness remains tunable (**Figure 3f**). Mechanical testing revealed the compressive modulus to be ∼29 kPa for soft fibers and ∼585 kPa for ‘stiff’ fibers, with the control group showing a similar modulus (∼575 kPa) (**Figure 3g**). Tensile testing further demonstrated that ‘stiff’ aligned (Stiff AL) scaffolds had the highest elastic modulus (∼52 MPa), nearly four times higher than other MeHA-dECM groups (**Figure 3h**).

These data confirm the successful fabrication of tendon dECM-based MeHA nanofiber scaffolds featuring key tendon ECM components and tunable mechanical properties, supporting their use in zone-specific tendon regeneration strategies.

### 2.3. Determining scaffold stiffness and alignment to enhance zone-specific regeneration in the rotator cuff enthesis

Next, we investigated how stiffness and alignment of the dECM-based scaffold system influenced cellular phenotype, aiming to identify optimal scaffold conditions for tendon (Phase 1) and fibrocartilaginous (Phase 2) regeneration in rotator cuff enthesis repair. To this end, bovine mesenchymal stem cells (bMSCs) or bovine tenocytes (bTCs) were cultured on the tunable dECM-based scaffolds.

The incorporation of tendon-derived dECM components within scaffolds significantly enhanced cell attachment, even in the absence of additional cell adhesion peptides such as RGD, compared to aligned 100% modified MeHA-only scaffolds (Ctrl), where cells appeared smaller, rounder, and less proliferative (**Figure S3a, Supporting Information**). bMSCs and bTCs cultured on the ‘stiff’ scaffolds were larger than those on the ‘soft’ scaffolds, and cells on aligned (AL) scaffolds exhibited higher aspect ratios than those on non-aligned (NAL) scaffolds, indicating the capacity of these scaffolds to regulate cell attachment and shape (**Figures 4a, b, c, Figures S3a, b, c, d, Supporting Information**). Cells exhibited high proliferation on all dECM-based scaffolds, indicating no scaffold-induced cytotoxicity (**Figure S3c, Supporting Information**).

**Figure 4.**
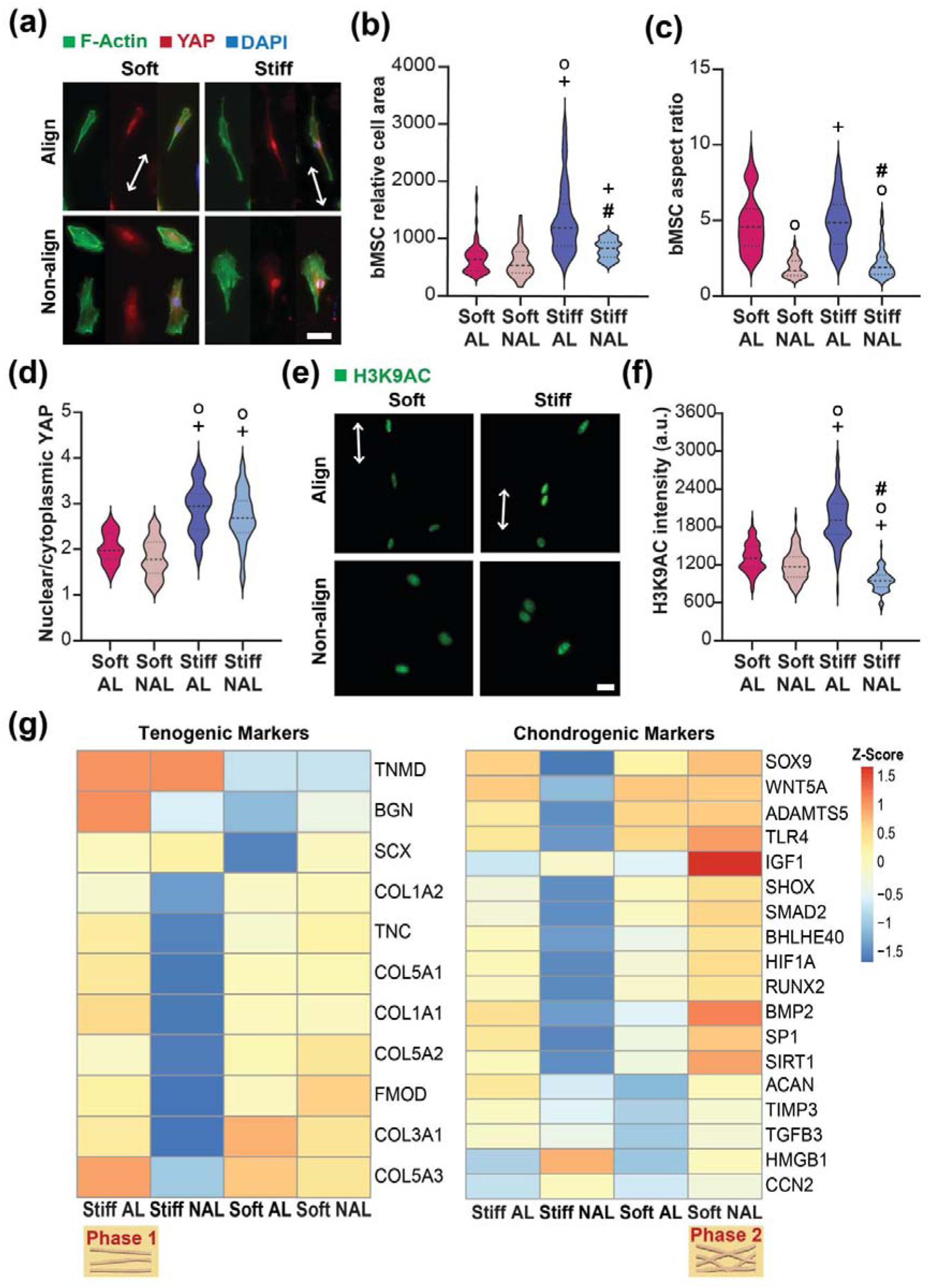
Tunable dECM scaffolds modulate cell morphology and gene expression. (a) Representative images of bMSCs cultured on dECM-based nanofibrous scaffolds for 3 days (Arrows: fiber direction; Green: F-actin, Red: YAP, Blue: DAPI; Align (AL), Non-align (NAL); Scale bar: 100 µm). Quantifications of relative (b) bMSC areas (n = 50/group; ○: p < 0.0001 vs. Soft AL, +: p < 0.01 vs. Soft NAL, #: p < 0.0001 vs. Stiff AL), (c) bMSC aspect ratios (n = 50/group; ○: p < 0.0001 vs. Soft AL, +: p < 0.0001 vs. Soft NAL, #: p < 0.0001 vs. Stiff AL), and (d) YAP nuclear localization ratios (n = 50/group; ○: p < 0.0001 vs. Soft AL, +: p < 0.0001 vs. Soft NAL, #: p < 0.0001 vs. Stiff AL). (e) Representative H3K9ac fluorescence images in bMSC nuclei in cells cultured on dECM-based scaffolds after 3 days (Arrows: fiber direction; Scale bar: 100 µm). (f) Quantification of H3K9ac fluorescence intensity in bMSC nuclei (n = 50/group; ○: p < 0.01 for Soft AL vs. Soft NAL, ○: p < 0.0001 Soft AL vs. Stiff AL,+: p < 0.0001 vs. Soft NAL, #: p < 0.0001 vs. Stiff AL). (g) Z-score heat maps of tenogenic gene expression via RNA-seq (n = 4/group from 4 biological replicates) and Z-score heat maps of chondrogenic gene expression via RNA-seq (n = 4/group from 4 biological replicates).

Furthermore, bMSCs seeded on ‘stiff’ nanofiber scaffolds demonstrated increased nuclear localization of YAP, a mechanosensitive transcriptional regulator, compared to those on soft scaffolds, regardless of alignment (**Figures 4a, d**). In addition, the fluorescence intensity of H3K9ac (Acetylation on histone H3 lysine 9, a marker for transcriptional activation ^[35]^) was significantly higher in the bMSC nuclei seeded on Stiff AL group compared to other dECM-contained scaffold groups (**Figures 4e, f**). These findings suggest that scaffold stiffness and alignment modulate epigenetic and transcriptional activity of cultured cells.

Indeed, RNA-seq analysis revealed that Stiff AL group enhanced the expression of tenogenic genes, indicating their potential suitability for promoting tendon zone (Phase 1) regeneration in rotator cuff enthesis repair (**Figure 4g**). In contrast, Soft NAL group promoted both chondrogenic and tenogenic gene expression, suggesting their potential for supporting regeneration of the fibrocartilaginous tendon-to-bone interface (Phase 2) (**Figures 4g)**. Additionally, RT-qPCR results showed that bTCs cultured on Stiff AL highly expressed fibrous and tenogenic genes, while those on Soft NAL scaffolds upregulated chondrogenic markers (**Figure S3e, Supporting Information**).

These data demonstrate that the stiffness and alignment of the dECM-based scaffold system regulate cell behavior and gene expression. The Stiff AL scaffolds exhibit a more pronounced tenogenic phenotype compared to other groups, making them well-suited for tendon regeneration (Phase 1), while the Soft NAL scaffolds enhance both chondrogenic and tenogenic phenotypes, making them appropriate for fibro-chondrogenic differentiation (Phase 2). Thus, these results indicate their potential to facilitate complex tissue integration within a biomimetic multiphasic scaffold system for rotator cuff enthesis repair.

### 2.4. Development of Bioactive POMaC/BG Composite Scaffolds for Enhancing Fibrocartilage-Bone Interface Regeneration

Since the fibrocartilaginous-to-bone region of the rotator cuff enthesis requires a biomaterial capable of supporting both mineralization and fibrocartilage-like tissue formation, we next developed a porous, composite scaffold for Phase 3. This phase is comprised of a citrate-based composite, POMaC, a bioenergetic molecule in bone physiology ^[26,27,29]^, and bioactive glass (BG), facilitating the exchange of network-modifier ions such as Na^+^ and Ca^2+^ with H^+ [31,32,34,36]^ (**Figure S1a, Supporting Information)**. To ensure robust integration between Phase 1/2 and Phase 3, the dECM-based nanofibrous scaffold was directly electrospun onto the POMaC-based porous scaffold. Free radical polymerization between MeHA and POMaC at their interface can create strong adhesion between the two phases (**Figure 5a**). Furthermore, hydrogen bonding between functional groups, including hydroxyl and carboxylic acid groups, present in both MeHA and POMaC further enhances this interfacial adhesion. Porous scaffolds containing 0 wt% (POMaC/0 BG) and 40 wt% BG (POMaC/40 BG) were fabricated via salt leaching, resulting in comparable pore sizes (355–500 μm) between groups (**Figure 5b, Figure S4a, Supporting Information**). Thermogravimetric analysis (TGA) confirmed approximately 40% BG incorporation in POMaC/40 BG (**Figure S4b, Supporting Information**). Fourier Transform Infrared Spectroscopy (FTIR) validated the interaction between POMaC and BG. Compared to POMaC/0 BG, POMaC/40 BG showed a distinct peak at 1599 cm^-1^ (**Figure 5c**), indicative of metallic carboxylate formation (sodium/calcium carboxylates or calcium dicarboxylate bridge) in the composite ^[37]^. This suggests that BG acts as a filler and a crosslinking agent within the composite. Incorporating BG into POMaC accelerated the degradation rate of the scaffolds, with 21.7% mass loss after 56 days in PBS (**Figure 5d**). In comparison, POMaC/0 BG exhibited less than 1% mass loss, suggesting that BG composites can modulate scaffold biodegradability. POMaC/0 BG and POMaC/40 BG scaffolds exhibited a three-phase swelling: an initial rapid absorption phase, followed by a slower swelling rate, culminating in a stable plateau (**Figure S4c, Supporting Information**). Notably, the POMaC/40 BG scaffold exhibited a lower swelling than POMaC/0 BG at Day 1 and Day 4, contributing to scaffold structural integrity (**Figure S4c, Supporting Information**).

**Figure 5.**
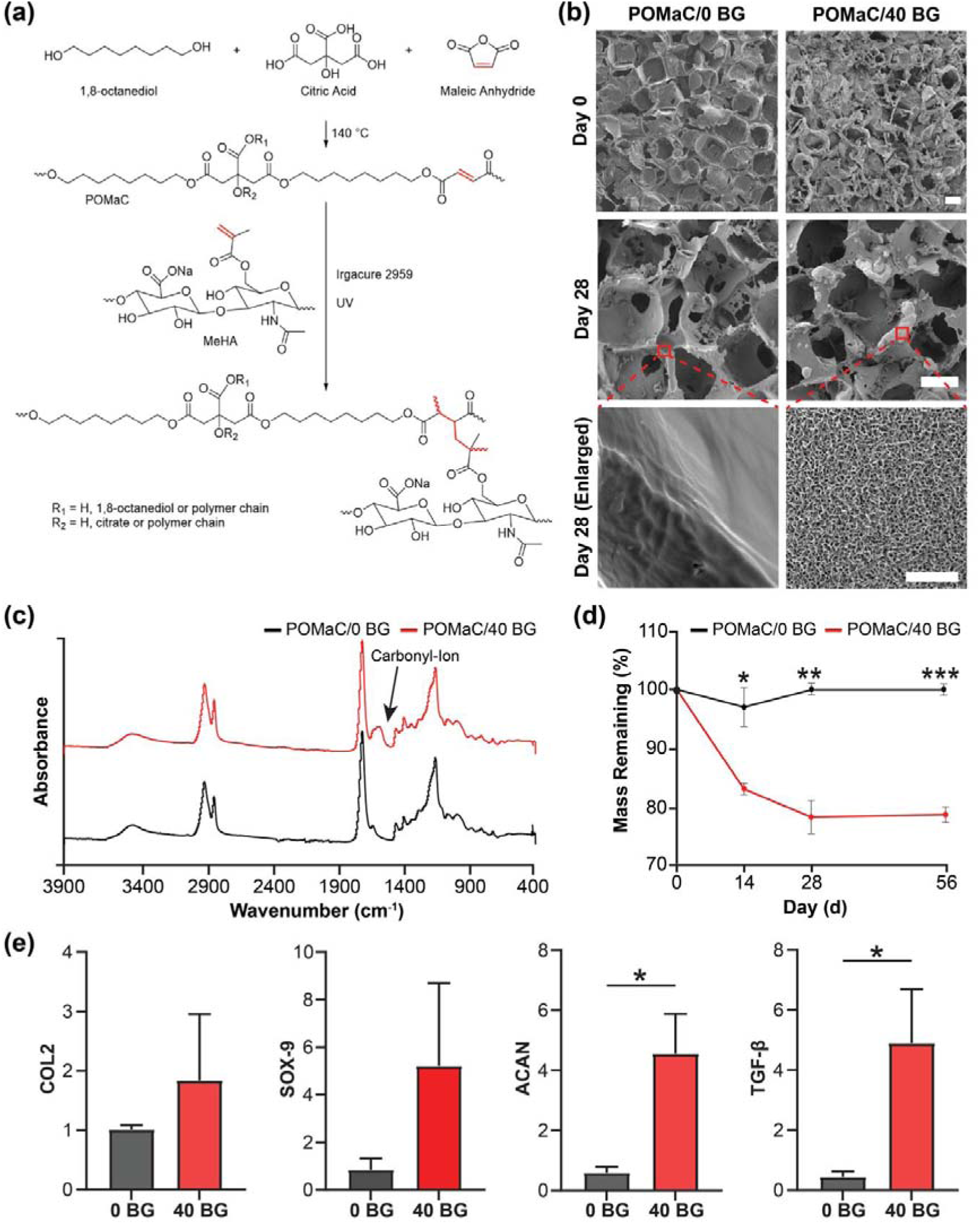
POMaC/Bioactive Glass Scaffolds for Fibrocartilaginous-Bone Interface Regeneration. (a) Schematic of radical polymerization between POMaC and MeHA forming a crosslinked network. (b) SEM images showing surface morphology and mineralization of POMaC/0 BG and POMaC/40 BG scaffolds on Day 0 and Day 28 (Scale bars: 200 µm for full images, 2 µm for zoomed-in images). (c) FTIR spectra of POMaC/0 BG and POMaC/40 BG scaffolds. (d) Degradation rates of both scaffolds in PBS over time (*: p < 0.05, **: p < 0.001, ***: p < 0.0001, n = 4/group). (e) Relative chondrogenic gene expression in bMSCs at Day 14 (*: p < 0.05, n = 5/group).

Assessing the ability of an implantable material to induce apatite formation in simulated body fluid (SBF) is crucial for predicting its *in vivo* bone interactions ^[38]^. Apatite, a calcium and phosphate-rich mineral, facilitates biomaterial bonding to native bone ^[39]^. A biomaterial is deemed bioactive if apatite precipitates on its surface within a defined period during SBF incubation. After 28 days in simulated body fluid (SBF), POMaC/0 BG scaffolds showed no obvious mineralization (**Figure 5b**, Day 28). In contrast, POMaC/40 BG scaffolds exhibited significant apatite formation, indicating enhanced bioactivity (**Figure 5b**, Day 28). Cultures of bMSCs on both POMaC/0 BG and POMaC/40 BG scaffolds exhibited significant cell proliferation, indicating no cytotoxicity from the scaffold components (**Figure S4d, Supporting Information**). Furthermore, bMSCs cultured on POMaC/40 BG showed significantly higher expression of chondrogenic markers [e.g., Collagen Type-2 (COL-2), SOX-9, Aggrecan (ACAN), and TGF-β compared to those on POMaC/0 BG, suggesting enhanced chondrogenic differentiation due to BG addition (**Figure 5e**).

These data indicate that integrating BG within POMaC scaffolds enhances mineralization, stability, and degradation. In addition, the elevated chondrogenic gene expression observed in POMaC/40 BG scaffolds indicates a microenvironment favorable for fibrocartilage-like differentiation, supporting its use within the calcified zone of the tendon (Phase 3) of the BMS. These findings suggest the potential of citrate-based biocomposite scaffolds to address the multifaceted requirements of enthesis regeneration by simultaneously promoting mineralization and creating an environment conducive to fibrocartilaginous tissue development.

### 2.5. Development and Evaluation of a Biomimetic Multiphasic Scaffold (BMS) for Zone- Specific Regeneration of the Rotator Cuff Enthesis

We identified the optimal scaffold compositions for each regenerative zone within the BMS by building on extensive *in vitro* evaluations of dECM-based nanofibrous scaffolds and POMaC-based porous scaffolds. Specifically, ‘stiff’ aligned dECM-based nanofibrous (Stiff AL) scaffolds were selected for Phase 1 (tendon regeneration), ‘soft’ non-aligned dECM-based nanofibrous scaffolds (Soft NAL) for Phase 2 (fibrocartilaginous regeneration), and POMaC/40 BG porous scaffolds for Phases 3 (fibrocartilage-bone regeneration) to promote the spatially stratified regeneration of the rotator cuff enthesis. To construct the BMS, Phase 1 and Phase 2 dECM-based nanofibrous structures were directly electrospun onto the POMaC/40 BG porous structure (**Figure 6a**). Picrosirius red and DTAF staining confirmed the structural integrity and stability of the BMS was maintained in basal media without phase separation (**Figure 6b**).

**Figure 6.**
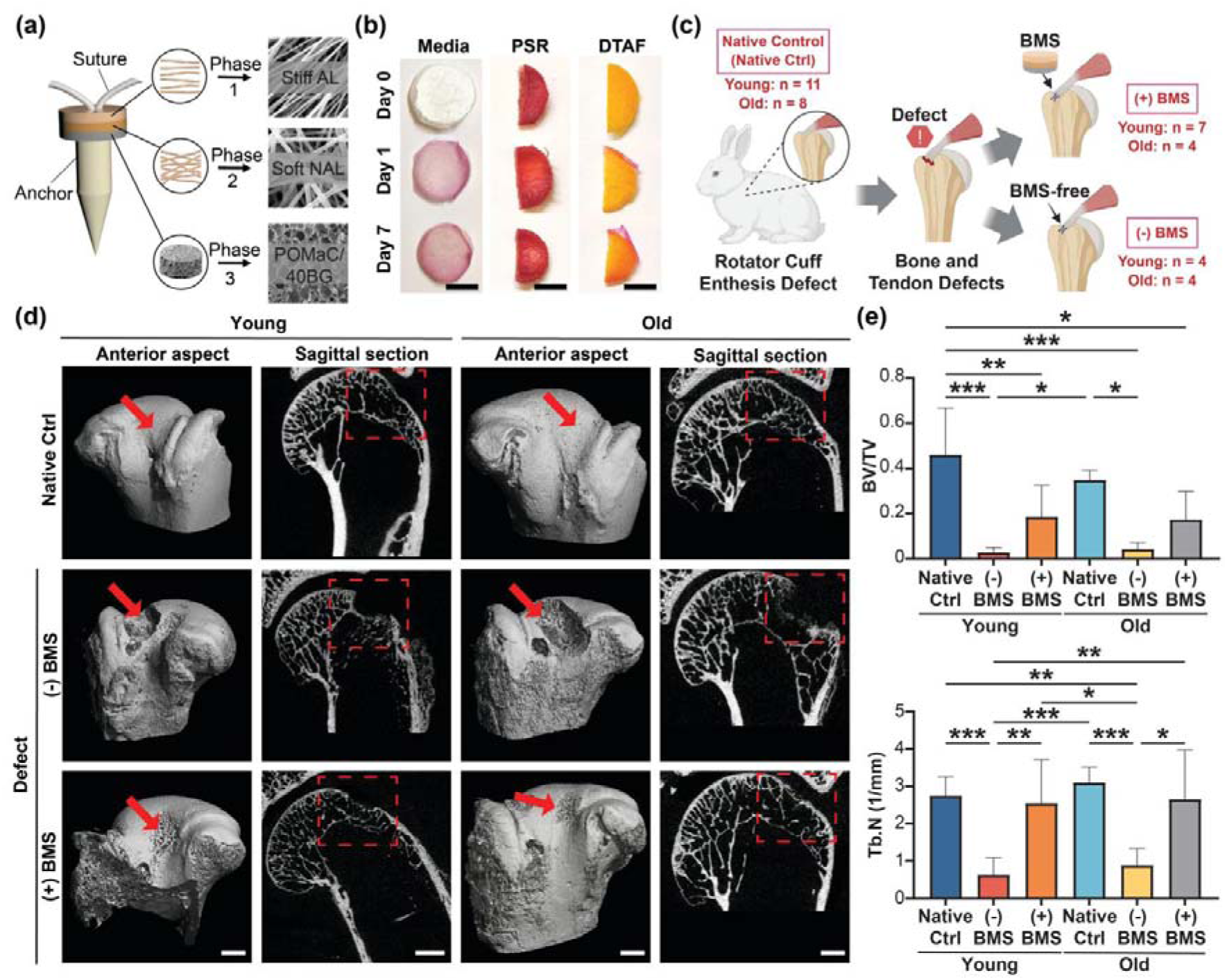
Micro-CT evaluation of the biomimetic multiphasic scaffold (BMS) in rotator cuff repair. (a) Schematic of BMS assembly illustrating the three-phase design for zone-specific regeneration of the rotator cuff enthesis. (b) Evaluation of structural stability and protein retention in the assembled BMS scaffold using Picrosirius Red and DTAF staining (Scale bar = 50 mm). (c) Schematic of the rabbit rotator cuff repair model and surgical procedure, including scaffold placement and experimental groups: Native Ctrl (native control; Young: n = 11, Old: n = 8), [(-) BMS] (defect alone; Young: n = 4, Old: n = 4), and [(+) BMS] (BMS treated; Young: n = 7, Old: n = 4). "Young" refers to rabbits <10 months old; "Old" refers to rabbits >3 years old. (d) Representative micro-CT images at 4 weeks post-surgery showing bone healing across Native Ctrl, [(-) BMS], and [(+) BMS] groups, with highlighted regions of interest. (e) Quantitative micro-CT analysis of bone remodeling, including bone volume/total volume (BV/TV) and trabecular number (Tb.N). Significant differences among groups are indicated (*: p < 0.005, **: p < 0.001, ***: p < 0.0001).

Next, to assess zone-specific tissue regeneration and integrative rotator cuff healing, the BMS was evaluated in proof-of-concept rotator cuff model in young and old female rabbits ^[40,41]^. BMS scaffolds were placed between the tendon and bone [BMS-treated group: (+) BMS] (**Figure 6c, Figure S5a, Supporting Information**). Control groups included untreated rabbit shoulders (Native Ctrl) and injured shoulders with only suture fixation, representing the defect group without BMS scaffolds [(-) BMS] (**Figure 6c**). The (+) BMS group included 7 young and 4 adult right shoulders; the (-) BMS group included 4 young and 4 adult right shoulders. Left shoulders served as native controls (**Figure 6c**).

All animals completed the study without adverse events. At 4 weeks post-operative, shoulders were harvested and processed for *ex vivo* analyses: micro-CT imaging revealed enhanced bone remodeling and regeneration in the (+) BMS groups compared to the (-) BMS groups (**Figure 6d, Figure S5b, Supporting Information**). Quantitative analysis showed significantly higher bone volume/total volume (BV/TV) and trabecular number (Tb.N) in the (+) BMS group, approaching values observed in native controls (**Figures 6e, f**). Additionally, trabecular separation (Tb.Sp) was significantly reduced in the (+) BMS group compared to the (-) BMS group (**Figure S5c, Supporting Information**). Notably, both young and adult rabbits treated with BMS scaffolds demonstrated similar improvements in bone regeneration, indicating that scaffold efficacy was consistent across age groups.

Next, to evaluate the integration of the biomimetic multiphasic scaffold with native rotator cuff tissues, histology using H&E, Masson’s Trichrome, Safranin O/Fast Green, and Picrosirius Red staining in young and old (-) BMS groups was characterized by disorganized tissue architecture, absence of fibrocartilage, and disrupted tendon-bone continuity. The insertion site exhibited immature tendon tissue formation and degenerated bone morphology, indicating poor repair (**Figures 7, 8**). In contrast, the (+) BMS groups showed substantially improved healing responses, with well-organized matrix formation and structural regeneration at the tendon-bone interface (**Figures 7, 8**). In particular, the old (+) BMS group also exhibited a well-defined gradient structure, with distinguishable tendon, fibrocartilage, and bone regions at the enthesis, which were absent in the corresponding (-) BMS groups **(Figure 8**). To quantitatively assess tissue regeneration, we applied a modified histological scoring system focused on collagen content, fiber organization, insertion continuity, and cellularity (**Table S1, Supporting Information)**. At 4 weeks, both young and old (+) BMS groups achieved significantly higher histological scores compared to their (-) BMS counterparts, approaching levels observed in native controls (**Figures 7b, c and Figures 8b, c**). These findings reflect enhanced collagen deposition, improved fiber alignment, greater enthesis insertion integrity, and increased cellular presence, indicating that BMS implantation promotes robust and zone-specific regeneration of the rotator cuff enthesis. Moreover, this approach offers a promising solution to overcome the limitations of conventional suture anchors, which lack the biological cues and structural guidance necessary for enthesis regeneration.

**Figure 7.**
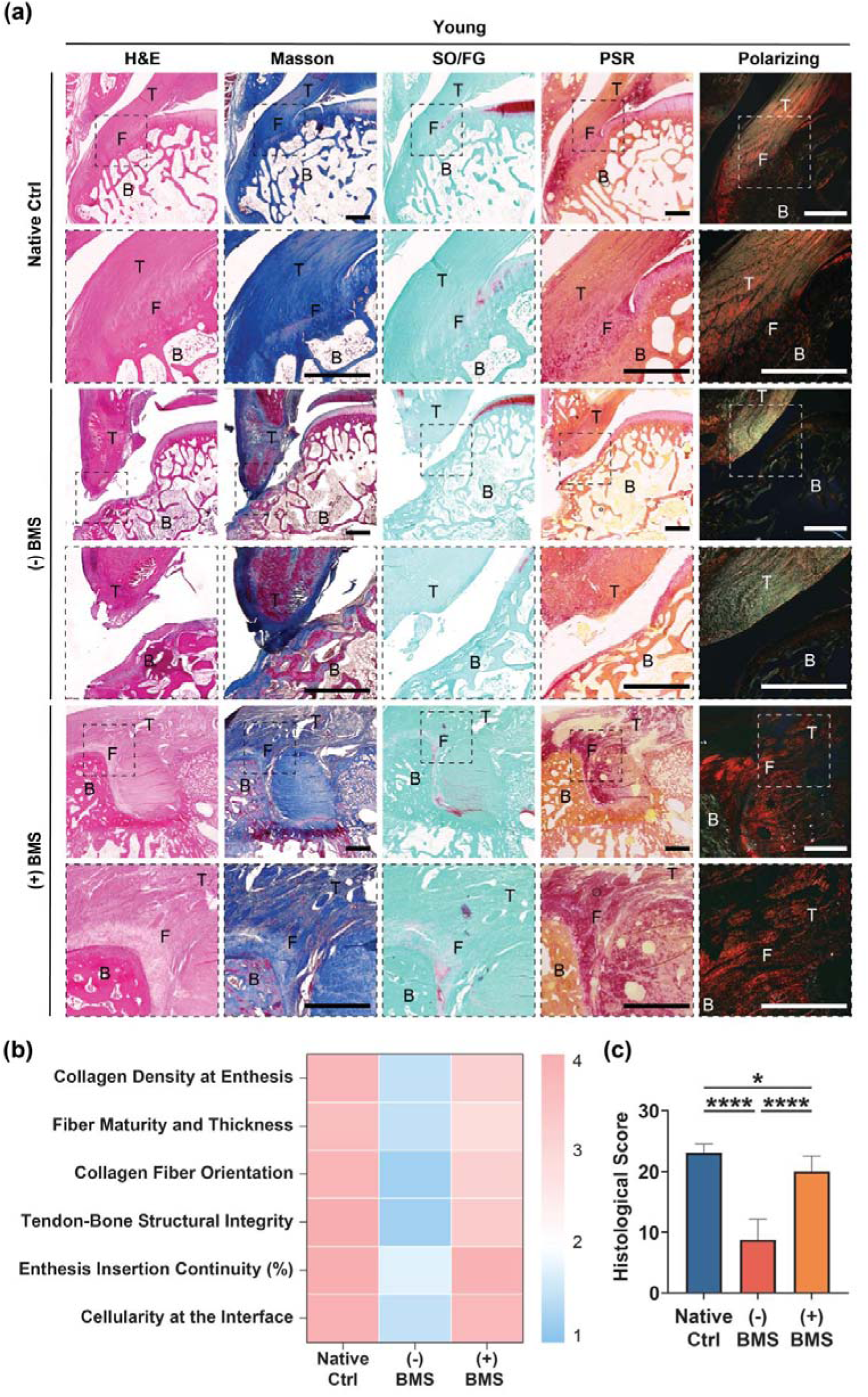
*In vivo* histological assessment of rotator cuff repair using the biomimetic multiphasic scaffold (BMS) in young rabbits. (a) Representative histological images of the tendon-to-bone interface at 4 weeks post-operation, stained with H&E, Masson’s trichrome, Safranin O/Fast green (SO/FG), and Picrosirius red (PSR), with corresponding polarized light micrographs. The (+) BMS group shows well-organized tissue architecture with distinct tendon, fibrocartilage, and bone layers, indicating successful integration and zonal regeneration in young rabbits (Scale bar = 1 mm; T = Tendon, FC = Fibrocartilage, B = Bone). (b) Heatmap summarizing histological scoring parameters across groups. (c) Quantitative analysis of total histological scores for tendon-enthesis units, demonstrating significantly improved outcomes in the (+) BMS group compared to controls (*: p < 0.005, ****: p < 0.00001).

**Figure 8.**
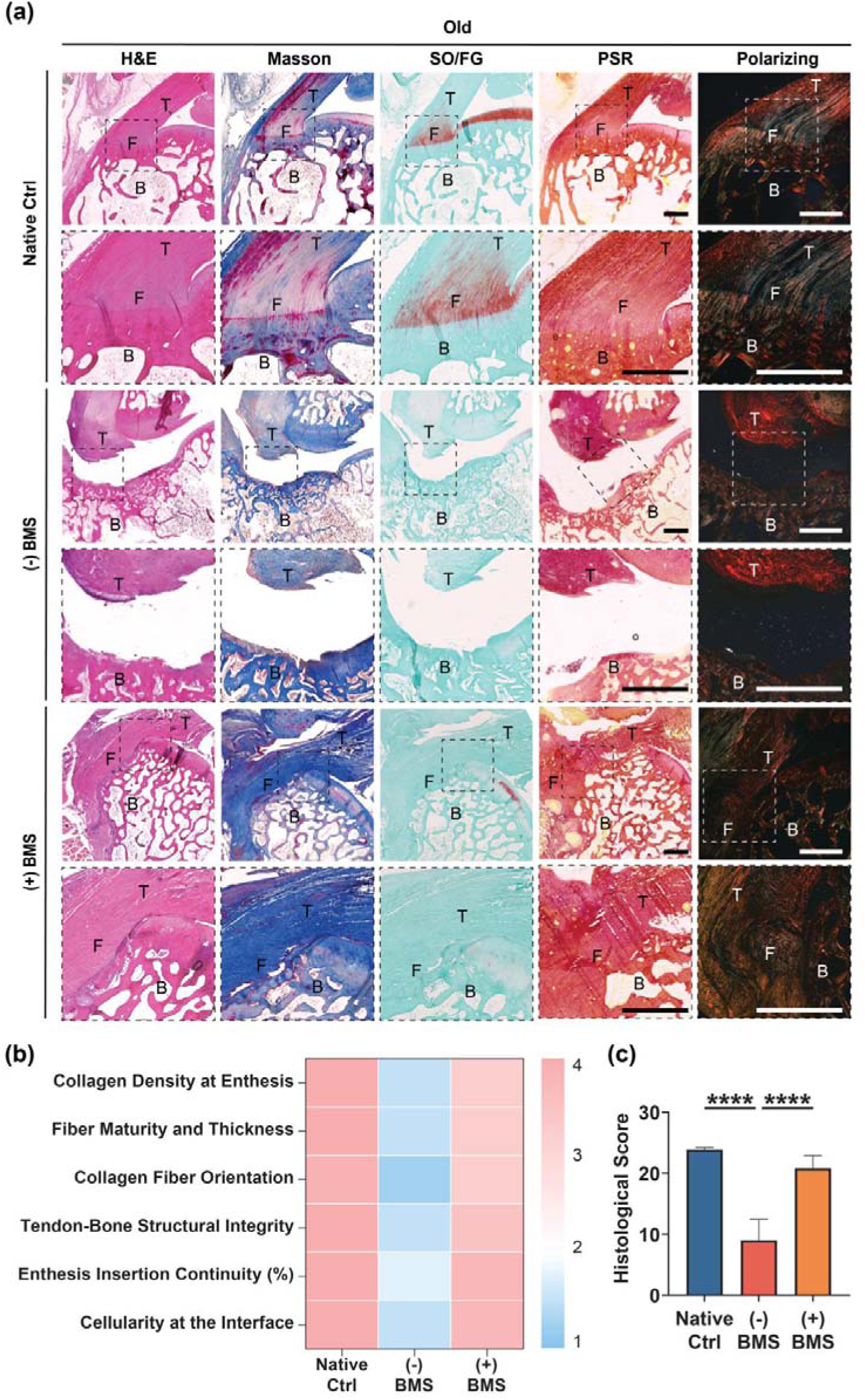
*In vivo* histological assessment of rotator cuff repair using the biomimetic multiphasic scaffold (BMS) in old rabbits. (a) Representative histological images of the tendon-to-bone interface at 4 weeks post-operation, stained with H&E, Masson’s trichrome, Safranin O/Fast green (SO/FG), and Picrosirius red (PSR), with corresponding polarizing light micrographs. The (+) BMS group shows well-organized tissue architecture with distinct tendon, fibrocartilage, and bone layers, indicating successful integration and zonal regeneration in old rabbits (Scale bar = 1 mm; T = Tendon, FC = Fibrocartilage, B = Bone). (b) Heatmap summarizing histological scoring parameters across groups. (c) Quantitative analysis of total histological scores for tendon-enthesis units, demonstrating significantly improved outcomes in the (+) BMS group compared to controls (****: p < 0.00001).

### 2.6. Discussion

Rotator cuff injuries are prevalent, and the gold standard repair method is suturing using hard polymer suture anchors ^[1,2,3]^. Unfortunately, re-tears occur frequently, with rates as high as 90% ^[8,9]^. This is often due to stress concentrations in these repaired regions ^[8,9]^. From a biomaterial perspective, commercial suture anchor systems focus solely on fixation to bone. However, the rotator cuff enthesis is a highly complex tissue that consists of roughly three phases: highly aligned tendon, a fibrocartilaginous intermediate region, and cartilage-bone ^[4,5]^. Thus, a new system that recapitulates the complex enthesis tissue structure would afford an enhanced healing process. To this end, this study addresses the critical limitations of current repair strategies by introducing a tissue-engineered multiphasic scaffold system that recapitulates the structural, mechanical, and biochemical properties of the native enthesis. This scaffold system provides biomimetic properties to current suture anchor systems, enhancing zonal development and regeneration, and potentially reducing re-tear rates.

To address these unmet clinical needs and overcome the existing method limitations, we developed a novel, tunable, biomimetic matrix-based multiphasic scaffold (BMS) through rigorous biological evaluations. In this study, our approach produced a ‘stiff’ aligned tendon decellularized tendon extracellular matrix (dECM)-based and ‘soft’ non-aligned dECM-based nanofibrous scaffold system for regenerating the tendon (Phase 1) and fibrocartilage (Phase 2) phases, respectively. In addition, a porous scaffold composed of poly(octamethylene maleate (anhydride) citrate) (POMaC) ^[26,27,28]^ and bioactive glass (BG) ^[30]^ facilitated regeneration of the fibrocartilage-bone phase (Phase 3). The BMS can be customized in size and delivered with traditional suture anchors, potentially providing essential bioactive and biophysical cues for effective enthesis tissue regeneration.

We selected decellularized bovine Achilles tendon as the base material, confirming that decellularization retained essential tendon matrix proteins, including collagen types I, II, III, V, and XI, biglycan, decorin, and fibromodulin. These proteins are pivotal for ECM structural support and organization, ensuring a conducive microenvironment for cellular adhesion, proliferation, and differentiation ^[42,43,44,45]^ in tendons. We combined this material with ‘stiff’ or ‘soft’ methacrylated hyaluronic acid (MeHA) ^[23,24]^ to create a tunable nanofibrous scaffold system using electrospinning and UV photocrosslinking. This maintained the dECM protein content while allowing for uncoupled stiffness and alignment modulation. The precise modulation of scaffold stiffness through MeHA incorporation is critical for mimicking the zonal architecture of the enthesis and regulating zone-dependent cell behavior and differentiation. *In vitro* studies demonstrated the ability of the BMS to regulate cell behavior and gene expression in a zone-dependent manner. The Stiff AL dECM scaffolds promoted tenogenic differentiation, which is evident from the increased expression of tendon-specific markers and activation of transcriptional regulators (e.g., enhanced YAP nuclear localization and H3K9ac intensity). Conversely, the Soft NAL scaffolds facilitated chondrogenic and tenogenic gene expression, supporting their role in regenerating the fibrocartilage interface. These findings indicate the importance of mechanical cues and ECM alignment in directing zone-specific cell differentiation and tissue formation within the rotator cuff enthesis. For enhanced fibrocartilage and bone integration (Phase 3), we developed a base structure of porous citrate-based material with enhanced osteogenic potential ^[26,28]^. Recent studies have shown reduced citrate levels in injured rotator cuff tissues, impairing essential cellular metabolism for tendon regeneration ^[11]^. Our citrate-based polymer, POMaC, addresses this metabolic deficit by locally supplying citrate to enhance cellular energetics, promote effective tendon-bone regeneration, and improve functional outcomes in rotator cuff repair ^[26]^. Bioactive glass (BG) ^[30]^ was incorporated into the POMaC scaffold to further enhance bone regeneration due to its ability to promote bioactivity and osteoconduction. The scaffold’s porous structure, particularly at a 40% BG concentration, enabled significant mineralization, critical for osteoconduction and effective bone regeneration.

Furthermore, the *in vivo* rabbit rotator cuff repair model demonstrated the translational promise of the BMS design. The *in vivo* tests confirmed that the scaffold facilitated integrative healing, with distinct zones of tendon, fibrocartilage, and bone forming at the repair site. Micro-CT imaging revealed enhanced bone remodeling at the fibrocartilage-to-bone interface as indicated by increased bone volume fraction (BV/TV) and trabecular number (Tb.N) alongside reduced trabecular separation (Tb.Sp). Thus, these findings suggest the (+) BMS group exhibited superior bone remodeling compared to the (-) BMS group featuring suture fixation alone. However, despite these improvements, the bone remodeling metrics in the (+) BMS group remained lower than those observed in the Native Ctrl, indicating that full restoration of native bone structure has yet to be achieved; given the relative short survival period, this is not surprising. Notably, the BMS performed similarly in both young and old donors, suggesting that its regenerative capabilities are not significantly impacted by donor age, making it applicable across a broad range of patients regardless of age-related tissue properties.

Histological analysis further revealed that implantation of the BMS significantly improved structural regeneration at the tendon-bone interface compared to suture-only repairs in both young and old rabbits. The clear demarcation of tendon, fibrocartilage, and bone zones in the BMS-treated tissues suggests that the scaffold’s spatially defined architecture and material composition supported zone-specific tissue remodeling. Particularly in older animals, where regenerative capacity is typically diminished, the BMS scaffold still promoted enthesis reconstruction with structural features resembling healthy native tissue. Although the in vivo results of this proof-of-concept study support the concept for the BMS in phase-distinct regeneration of the enthesis, this rabbit model represents an acute repair environment with assessments conducted at four weeks post-operative. Clinical rotator cuff patients often present with a history of chronic injury and re-injury. Thus, extending the postoperative survival time combined with a chronic rotator cuff animal model will be an essential next step to continue to develop this technology using translational test systems of high fidelity and rigor. Importantly, the BMS’s long-term durability and mechanical performance under physiological loading conditions need further evaluation. Additionally, scaling up the BMS for larger animal models is crucial for validating its translatable potential. Future research should also investigate incorporating bioactive molecules, such as growth factors or small biomolecules, to further enhance the scaffold’s regenerative capacity.

## 3. Conclusion

In conclusion, this study presents a promising multiphasic scaffold system that preserves tendon-derived bioactive proteins and elicits favorable changes in cellular behavior. The scaffold system significantly enhanced the tendon-bone interface in a proof-of-concept rabbit model, effectively mimicking the histological characteristics of native tissue. This biomimetic multiphasic scaffold system may address the limitations of current repair strategies using suture anchors alone and demonstrates significant potential for zone-specific tissue regeneration. Beyond rotator cuff applications, the principles shown in this study could be extended to other orthopedic interfaces, such as ligament and meniscus repair. The modularity of the BMS design allows for customization for various tissue interfaces, broadening its potential impact in musculoskeletal soft tissue regeneration.

## 4. Experimental Section

### Decellularized bovine Achilles tendon ECM preparation

The decellularized bovine ECM (dECM) matrix was prepared according to a previously established protocol ^[22]^. Briefly, juvenile bovine Achilles tendon tissue was harvested and cut into small cubes, each with a side length of 1 mm. These tissue cubes were stirred and incubated in a solution of 1% SDS/ phosphate-buffered saline (PBS) (w/v) (Invitrogen, USA; Gibco, USA) for 3 days at room temperature, with the solution being refreshed daily. The tissue samples were then washed with 0.1% EDTA/PBS (w/v) (Thomas Scientific, USA; Gibco, USA) solution for 1 day at room temperature. After an overnight rinse in deionized water (DI water) at room temperature, the tissue samples were further lyophilized in a freezer dryer (Labconco, USA) for 3 days and stored frozen at -20 °C for long-term preservation.

### MeHA synthesis

Methacrylated hyaluronic acid (MeHA) was synthesized according to published protocols, achieving modification degrees of either 35% (for ‘soft’) or 100% (for ‘stiff”) ^[23,46]^ as described previously. Sodium hyaluronate (60 kDa, Lifecore, USA) was dissolved in DI water overnight and reacted with calculated methacrylic anhydride (Sigma Aldrich, USA) on ice for 1.5 days. The pH of the solution was maintained between 8.5 and 9.5 by adding 5M NaOH (Sigma Aldrich, USA) dropwise. The MeHA solution was dialyzed in DI water at room temperature for 14 days and then frozen at −80 °C for 3 hours. Finally, the MeHA was lyophilized in a freezer dryer (Labconco, USA) for 3 days. The freeze-dried dECM was ground into powder and stored at -20 °C for subsequent experiments.

### Proteomic analysis

Lyophilized native and dECM bovine tissue powders were submitted to the Proteomics Core Facility at the Children’s Hospital of Philadelphia Research Institute for liquid chromatography-tandem mass spectrometry (LC-MS/MS) analysis ^[46]^. Data processing and statistical analysis were conducted using the Spectronaut™ Pulsar and Perseus software platforms (Max Planck Institute, Germany). To identify ECM components, the Gene Ontology term (GO:0031012 Extracellular Matrix) specific to Bos taurus organism was used, along with the MatrisomeDB1 ECM-protein knowledge database to delineate the core matrisome preserved in the dECM group. Protein-protein interaction network of the preserved core matrisome in dECM was constructed using STRING web tool (version 10.0, String Consortium 2020) and visualized with Cytoscape software (version 3.10.3). Gene Ontology analysis was performed using gProfiler, with biological processes, cellular components, and molecular functions classified as significant with Benjamini-Hochberg corrected p-value < 0.05. For visualization, stacked bar plots and bubble plots were generated using the ggplot2 package in R programming ^[47]^.

### Fabrication of dECM-based nanofibrous scaffolds

The ‘stiff’ aligned dECM-based nanofibrous scaffold electrospinning solution was prepared from 4% (w/v) ‘stiff’ MeHA, 2% (w/v) dECM, 2% (w/v) poly(ethylene oxide) (PEO, 900kDa, Acros Organics, USA), and 0.5% (w/v) Irgacure 2959 (Sigma Aldrich, USA) in DI water ^[22,23]^. The ‘soft’ nonaligned dECM-based nanofibrous scaffold electrospinning solution was prepared from 4% (w/v) ‘soft’ MeHA, 2% (w/v) dECM, 2% (w/v) PEO, and 0.5% (w/v) Irgacure 2959 in DI water ^[22,23]^. Phase 1 and Phase 2 scaffolds were fabricated by electrospinning using an 18-gauge metal needle (Hamilton, USA) with a voltage of 17 kV onto a grounded collector with a voltage of – 3 kV ^[22,23]^. The distance between the needle and collector was 15 cm and the flow rate was 1 mL h^-1^. Scaffolds were purged with nitrogen and photo-crosslinked with UV light (1.7 mW cm^-2^, 365 nm, UVP/Analytik Jena) for 20 minutes.

### Fabrication of 3D porous POMaC/BG scaffolds

Poly(octamethylene maleate citrate) (POMaC) was synthesized via condensation polymerization of 1,8-octanediol (Sigma Aldrich, USA), citric acid (Sigma Aldrich, USA), and maleic anhydride (Sigma Aldrich, USA) at 140°C under a nitrogen atmosphere for 3 hours, yielding pre-POMaC containing vinyl groups and ester bonds for further crosslinking ^[27]^. POMaC was covalently crosslinked with MeHA through free radical polymerization. Briefly, the polymers were combined with a photoinitiator (Irgacure 2959) and exposed to UV light to generate free radicals, triggering double bond cleavage and subsequent polymer crosslinking. This reaction resulted in the formation of a hybrid, crosslinked polymer network. Phase 3 scaffolds were composed of POMaC and bioactive glass (BG, 0/40 wt%, Mo-Sci Corporation, USA). The resulting POMaC pre-polymer was dissolved in 1,4-dioxane (Sigma Aldrich, USA) and mixed with BG (0/40 wt%, Mo-Sci Corporation, USA) and sodium chloride (92 wt%; Fisher Scientific, USA). The mixture was then molded and thermally crosslinked at 80 °C for 7 days. After crosslinking, the scaffolds were immersed in DI water to remove the sodium chloride and subsequently dried by lyophilization for future use.

### Scaffold characterizations

dECM-based nanofibrous scaffolds and POMaC/BG scaffold were coated with iridium by a sputter coater (Quorum Q150T ES, UK). Scanning electron microscope (SEM, FEI Quanta, USA) was used to characterize the scaffold morphology and porosity. Environmental scanning electron microscopy (ESEM, TFS Quanta 600, USA) was used to observe the hydrated nanofibers. To evaluate the mineralization capacity of the scaffold surface, POMaC/0 BG and POMaC/40 BG scaffolds were immersed into a simulated body fluid (SBF), designed to mimic the human blood plasma ^[48]^, for up to 28 days. To assess protein preservation, 100 µm thick sections of native bovine ECM, bovine dECM, and dECM-based nanofibrous scaffold were stained with Hematoxylin and Eosin (H&E) for nuclei visualization, Alcian Blue for glycosaminoglycan (GAG) retention, Picrosirius Red for collagen distribution, and 5-(4,6-dichlorotriazinyl) aminofluorescein (DTAF, Sigma Aldrich, USA) for overall protein detection ^[22,49]^. A Nikon widefield microscope (Nikon Instruments, USA) was used to visualize and capture images of nuclei, GAG preservation, and collagen distribution across the samples.

### Mechanical testing

Uniaxial tensile tests were performed on ‘soft’ and ‘stiff’ dECM/MeHA scaffold strips (width = 5 mm, length = 30 mm) using the Instron 5542 with a loading stress rate of 0.2%/second. Scaffold thickness was measured using a customized system described in a previous study ^[22]^. The elastic modulus values were calculated based on the linear region of the stress-strain curve obtained during the test. Compressive moduli of the nanofibrous scaffolds were determined by atomic force microscopy (AFM, Asylum MFP-3D, USA). Unconfined compression tests were performed on porous scaffolds using an eXpert 8613 mechanical tester (ADMET Inc., USA). Briefly, the scaffolds were compressed at a rate of 1.3 mm/min to 25% of the initial length. The initial modulus was calculated by measuring the gradient of the stress-strain curve at 10% compression.

### Fourier-transform infrared spectroscopy and accelerated degradation testing

Fourier-transform infrared spectroscopy (FT-IR) was acquired on a Thermo Electron Corp Nicolet 380 FT-IR Spectrometer (Thermo Fisher Scientific, USA) with 64 scans. The spectra were normalized at 1723 cm-1 for a better comparison. The accelerated degradation rate of porous scaffolds was assessed *in vitro* in PBS at 57°C for up to 56 days under static conditions. PBS was changed every week to maintain the pH above 7. At the predetermined time point, the samples were washed with DI water and lyophilized. The weight loss was calculated by comparing the initial weight (W_0_) with the weight measured at 0, 14, 28, and 56 days (W_t_). The degradation percentage was calculated using the following formula.

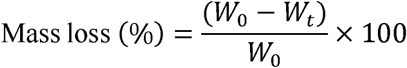

### Cell isolation and culture

Juvenile bovine knee joints (2–3 months old) were purchased (Research 87, USA). Bovine tenocytes (TCs) and bone marrow mesenchymal stem cells (bMSCs) were isolated from juvenile bovine Achilles tendons and tibia bone marrow, respectively. The basal cell culture medium was prepared using highglucose Dulbecco’s minimal essential medium (DMEM, Gibco, USA) with 10% fetal bovine serum (FBS, R&D systems, USA) and 1% Penicillin Streptomycin solution (Corning, USA). Passage 1 cells were collected and seeded on the scaffolds after reaching 90% confluency rate.

### Immunofluorescence staining

After 3 days of culture on dECM-based nanofibrous scaffolds, passage 1 bMSCs were fixed with 4% paraformaldehyde at room temperature and blocked with 1% bovine serum albumin (BSA, Sigma Aldrich, USA). To assess YAP nuclear localization, samples were incubated with anti-YAP mouse monoclonal IgG2a antibody (1:200, Santa Cruz Biotechnology, USA) followed by Alexa Fluor-546 goat anti-mouse IgG antibody (1:200, Life Technologies Corporation, USA). The cytoskeleton was stained with Alexa Fluor-488 phalloidin (Life Technologies Corporation, USA) at 37°C for 1 hour, and cell nuclei were stained by 4,6-diamidino-2-phenylindole (DAPI, Invitrogen, USA). For transcriptional activation analysis, Acetyl-H3K9 (H3K9ac) primary antibody (Invitrogen, USA) was applied, followed by Alexa Fluor 488 goat anti-rabbit secondary antibody (Invitrogen, USA) ^[50]^. Imaging was performed using a Leica fluorescence microscope, and cell morphology was quantified with Fiji software.

### Cell proliferation and gene expression on scaffolds

Passage 1 bTCs or bMSCs were seeded at 5,000 cells per scaffold (n = 5/group). Cell proliferation was assessed by adding 440 µL of Cell Counting Kit-8 reagent (1:10 dilution, Dojindo, Japan) to each scaffold and incubating at 37°C for 4 hours under 5% CO□. Absorbance was measured at 450 nm using a BioTek Synergy H1 microreader (Agient, USA). mRNA was isolated from cells utilizing the TRIzol (Fisher Scientific)/chloroform (Fisher Scientific) extraction method ^[46]^. Total RNA concentrations were measured using a NanoDrop spectrophotometer (NanoDrop Technologies, USA) ^[51]^. Complementary DNA (cDNA) was synthesized using the PhotoScript II First-Strand cDNA Synthesis Kit (New England BioLabs, USA). Quantitative reverse transcription polymerase chain reaction (RT-qPCR) was performed using a QuantStudio 6 Pro (Applied Biosystems, USA) and Fast SYBR Green Master Mix (Applied Biosystems, USA). The expression levels of collagen type I alpha 2 chain (COL1A2), collagen type II (COL2), collagen type III (COL3), tenascin-C (TNC), tenomodulin (TNMD), and aggrecan (ACAN) were measured. Expression levels were normalized with respect to the housekeeping gene glyceraldehyde 3-phosphate dehydrogenase (GAPDH). Primer sequences are listed in **Table S2** (**Supporting Information**).

### RNA sequencing

RNA sequencing was conducted on Passage 1 bovine mesenchymal stem cells (MSCs) cultured on dECM/MeHA nanofibers with different stiffness and alignment configurations: Stiff AL, Stiff NAL, Soft AL, and Soft NAL. Total RNA was extracted using the TRIzol (Fisher Scientific, USA)/chloroform (Fisher Scientific, USA) method. RNA-Seq library preparation, including ribosomal RNA (rRNA) depletion and 150 bp paired-end sequencing, was performed at Genewiz (South Plainfield, USA) ^[46]^. Sequence reads were processed with Trimmomatic v.0.36 for adapter and low-quality nucleotide removal, then aligned to the Bos taurus reference genome (Ensembl) using the STAR aligner v.2.5.2b ^[46]^. Unique gene hit counts were generated via FeatureCounts from the Subread package v.1.5.2. Differential expression analysis was performed in R using DESeq2 ^[46]^. Differential gene expression was assessed using a Likelihood Ratio Test (LRT), with statistical significance defined as p < 0.05 ^[46]^. Differentially expressed genes (DEGs) were identified based on an adjusted p-value < 0.05 (Benjamini-Hochberg) and |log□(fold change)| > 1.0 ^[46]^.

### Rabbit rotator cuff in vivo study design

BMS was assembled by electrospinning ‘stiff’ aligned dECM-based nanofibrous scaffold (Stiff AL) for 6 hours and ‘soft’ nonaligned dECM-based nanofibrous scaffold (Soft NAL) for another 6 hours onto POMaC/ 40 BG scaffold with 20-minute UV light cross-linking (1.7 mW cm^-2^, 365 nm, UVP/Analytik Jena, USA). All *in vivo* procedures were approved by the University of Pennsylvania Institutional Animal Care and Use Committee (IACUC). Each rabbit’s left shoulder served as the control (Native ctrl), while the right shoulder underwent either BMS implantation [(+) BMS] or negative control [(-) BMS] surgical treatment. Rabbits were divided into two age groups: Young (<10 months old) and Old (>3 years old). For the (+) BMS treatment, the right shoulders of 7 young and 4 old rabbits were utilized, while the (-) BMS treatment was applied to the right shoulders of 4 young and 4 old rabbits. The left shoulders of all rabbits were collected as part of the positive control group.

This study was reviewed and approved by the Institutional Animal Care and Use Committee of the University of Pennsylvania. Skeletally mature, female, New Zealand White rabbits were enrolled in the study. The animals were sourced from a commercial supplier (Charles River Laboratories Inc., USA). Animals received a subcutaneous injection of pre-anesthetic and general anesthesia consisting of Ketamine 35mg kg^-1^ and Xylazine 5mg kg^-1^. Following endotracheal intubation, animals were maintained under Isoflurane/oxygen for the duration of the procedure. Using aseptic technique, the right supraspinatus muscle and tendon served as the experimental side in all animals, with the left shoulder as an unoperated control. Briefly, an open craniolateral approach was performed on the right shoulder, approaching the supraspinatus muscle via splitting of the deltoid muscle followed by sharp transection of the right supraspinatus tendon from the insertion at the greater tuberosity. Two bone tunnels were drilled on both the supraglenoid and greater tubercle using 1mm Kirschner wires. Bone troughs perpendicular to the direction of the supraspinatus muscle fiber on both the medial margin of the greater tubercle and the dorsal aspect of the glenoid were created. The tendon of the supraspinatus was secured using a modified Mason-Allen suture enhancement technique, followed by passing two strands each of #1 PDS suture (Ethicon, USA) through each bone tunnel and tying at the level of the greater tubercle. One strand of suture of each bone tunnel was knotted in a parallel pattern. The remaining strand of suture of one bone tunnel was knotted with the strand of another bone tunnel in a crossed pattern. BMS scaffolds were placed between the tendon and bone [BMS-treated group: (+) BMS]. Control groups included untreated rabbit shoulders (Native Ctrl) and injured shoulders with only suture fixation, representing the defect group without BMS scaffolds [(-) BMS]. The (+) BMS group included 7 young and 4 adult right shoulders; the (-) BMS group included 4 young and 4 adult right shoulders. The left shoulders served as native controls. The deltoid and skin incision were closed in layers with 2-0 Monocryl sutures (Ethicon, USA). Postoperatively, animals were allowed free cage activity. During the postoperative time, clinical parameters including appetite, activity, signs of infection, bleeding, and wound dehiscence were evaluated daily by a veterinary surgeon. At 4 weeks post-operative, animals were euthanized with a commercially available euthanasia solution (Pentobarbital 1mL 5kg^-1^) according to the guidelines set forth by the current AVMA Panel on Euthanasia and shoulders were harvested and processed for ex vivo analyses.

### Micro-CT scanning

Rabbit shoulders were fixed in 10% neutral buffered formalin (NBF, Sigma Aldrich, USA) for 4 days and then subjected to ex vivo μCT scanning using a Scanco μCT45 system (Scanco Medical AG, Switzerland). Scans were performed at an 18.5-μm voxel resolution with a 300 ms integration time, 145 μA current, and 55 kVp energy. A 2 mm thick region of interest (ROI) was selected from μCT images of each humerus bone to evaluate bone volume fraction (BV/TV), trabecular thickness (Tb.Th), and trabecular separation (Tb.sp).

### Histological analysis

After micro-CT scanning, rabbit shoulders were decalcified at room temperature using Formical-2000 for 3 weeks. After decalcification, the samples were dehydrated using graded ethanol (Decon Labs, Switzerland), embedded in paraffin (Azer Scientific, USA) and sectioned into slices with a thickness of 8 μm by a rotary microtome (Leica, Germany). After sectioning, histological staining was conducted using Hematoxylin and Eosin, Masson’s Trichrome, Safranin O/Fast Green, and Picrosirius Red staining ^[22,46]^. Polarizing light microscopy was carried out using a Leica DMLP Polarizing Light Microscope (Leica Microsystems, Germany). The stained slices were imaged using a Zeiss Axiocam 506 mono camera (Zeiss, Germany), and images were processed with Zeiss ZEN 3.9 software (Zeiss, Germany).

### Statistical analysis

Experimental results were statistically analyzed using GraphPad Prism 9 (GraphPad Software, Inc., USA). Student’s t-test or one-way analysis of variance (ANOVA) with Tukey’s post hoc test was employed to determine significant differences. Data are presented as mean ± standard deviation (SD), with significance defined as p < 0.05.

## Acknowledgements

The authors sincerely thank Wen Sang and Edgardo Arroyo for their support throughout this research. We also thank Shreya Mondal, Lucy Hederick, Erik Nimmer, Jason A. Burdick, and Robert L. Mauck for their scientific assistance and Penn Vet New Bolton Center staff for help with animal care. Micro-CT scan was performed in Penn Center for Musculoskeletal Disorders (PCMD) MicroCT Imaging Core (NIH/NIAMS P30-AR069619).

## Data Availability Statement

The data that support the findings of this study are available from the corresponding author upon reasonable request.

## Supporting Information

### Supplementary Materials and Methods

#### Chemicals

SDS, 4,6-diamidino-2-phenylindole (DAPI), Acetyl-H3K9 (H3K9ac) primary antibody, and Alexa Fluor 488 goat anti-rabbit secondary antibody were purchased from Invitrogen. phosphate-buffered saline (PBS), Dulbecco’s minimal essential medium (DMEM) were purchased from Gibco. EDTA was purchased from Thomax Scientific. 60 kDa Sodium hyaluronate was purchased from Lifecore. Methacrylic anhydride, 5M NaOH, Irgacure 2959, 1,8-octanediol, citric acid, maleic anhydride, 1,4-dioxane, 5-(4,6-dichlorotriazinyl) aminofluorescein (DTAF), bovine serum albumin (BSA), and neutral buffered formalin (NBF) were purchased from Sigma Aldrich. 900 kDa poly(ethylene oxide) (PEO) was purchased from Acros Organics. Bioactive glass (BG) was purchased from Mo-Sci Corporation. Sodium chloride, TRIzol, and choloroform were purchased from Fisher Scientific. Fetal bovine serum (FBS) was purchased from R&D systems. Penicillin Streptomycin solution was purchased from Corning. Anti-YAP mouse monoclonal IgG2a antibody was purchased from Santa Cruz Biotechnology. Alexa Fluor-546 goat anti-mouse IgG antibody and Alexa Fluor-488 phalloidin were purchased from Life Technologies Corporation. Cell Counting Kit-8 reagent was purchased from Dojindo. PhotoScript II First-Strand cDNA Synthesis Kit was purchased from New England BioLabs. Fast SYBR Green Master Mix was purchased from Applied biosystems. Graded ethanol was purchased from Decon Labs. Paraffin was purchased from Azer Scientific.

### Supplementary Figures and Legends

**Supplementary Figure S1.**
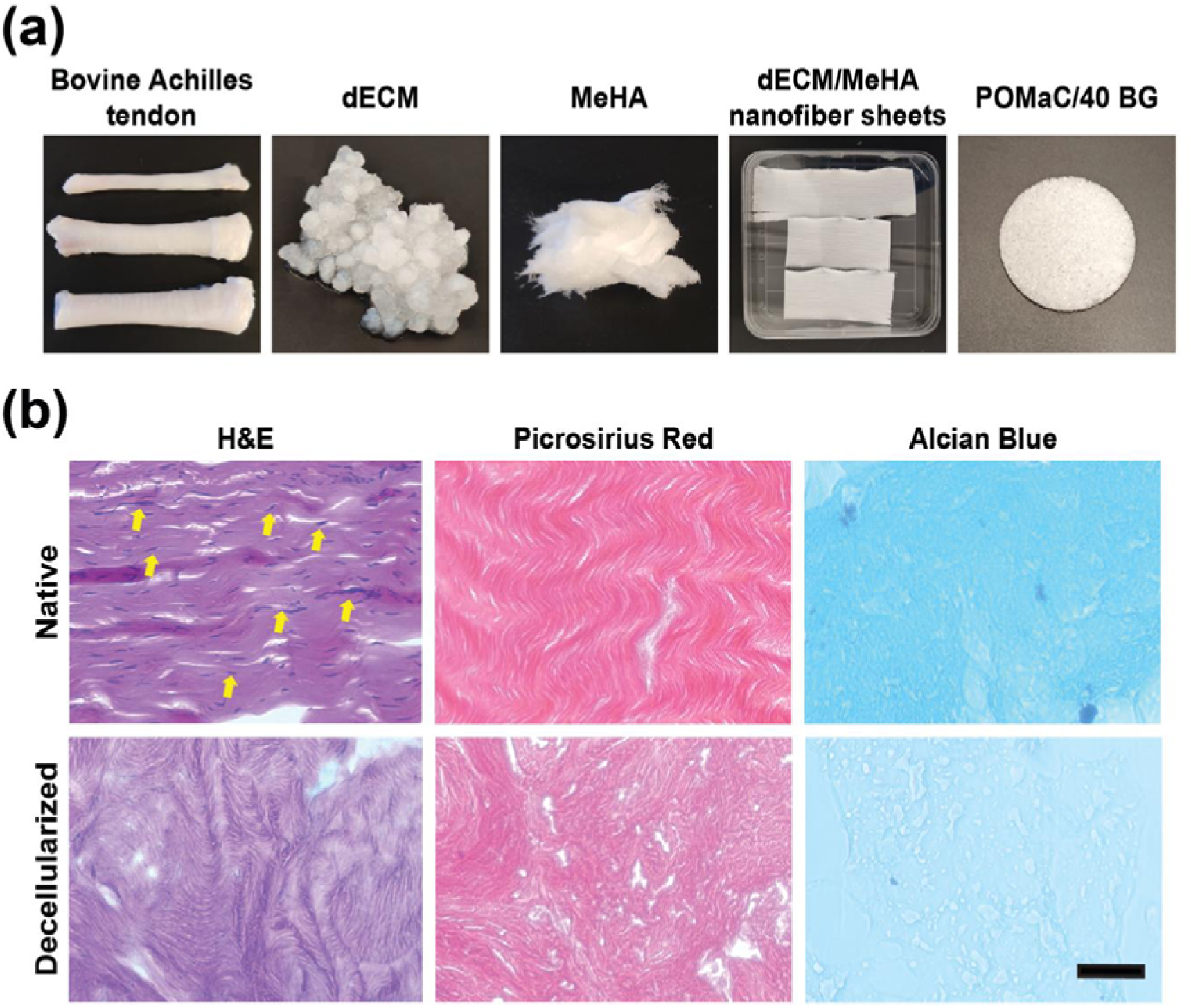
(a) Representative images of bovine Achilles tendon, decellularized ECM (dECM), stiff MeHA (modification rate = 100%), electrospun dECM/MeHA nanofiber scaffolds (top: stiff aligned (AL); middle: soft non-aligned (NAL); bottom: soft non-aligned (NAL)), and porous POMaC/40% BG scaffold. (b) Representative H&E, Picrosirius Red, and Alcian Blue staining of native and decellularized ECM tissues (Arrows indicate nuclei, scale bar = 50 µm).

**Supplementary Figure S2.**
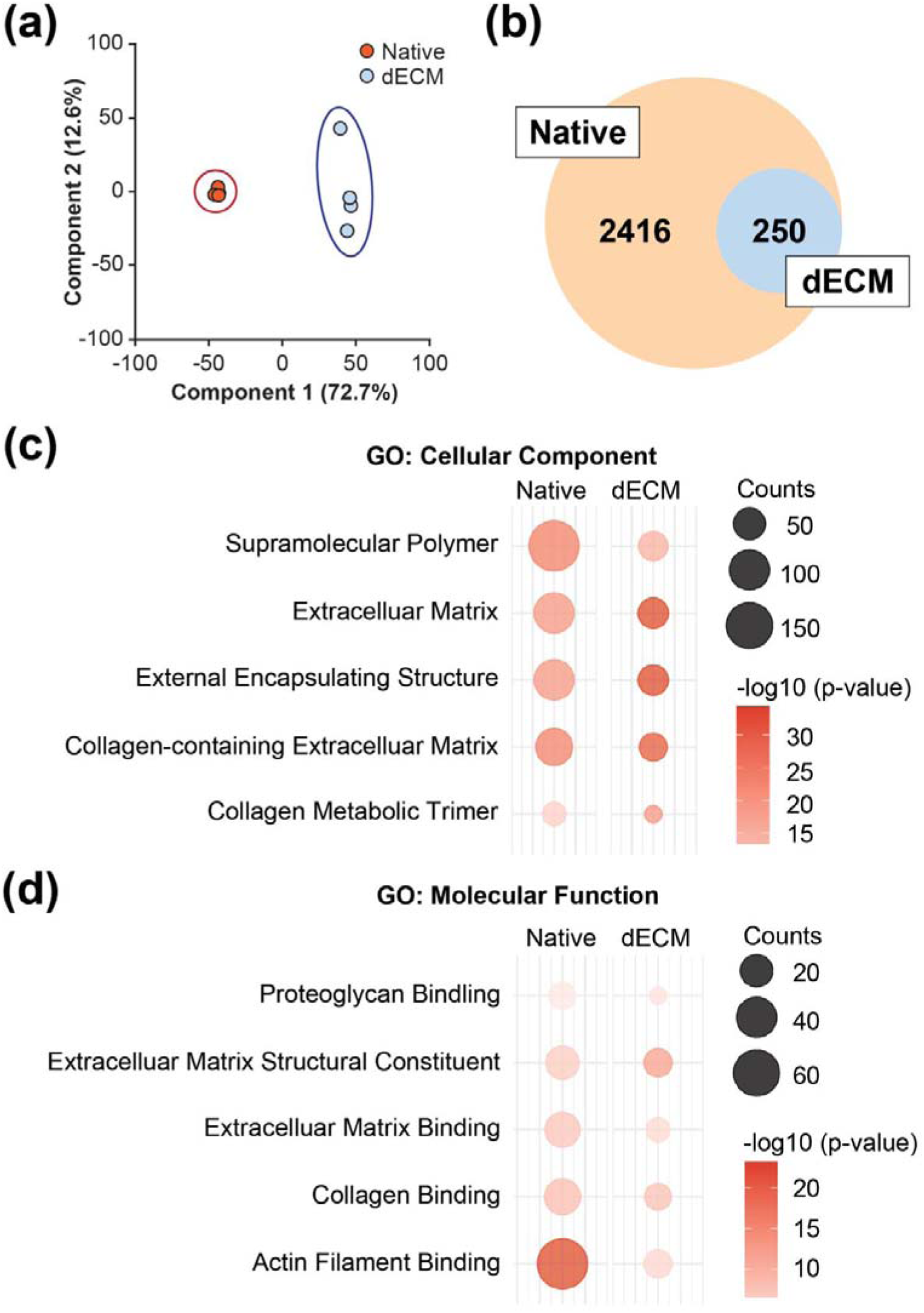
(a) Principal component analysis (PCA) showing distinct clustering of native and dECM samples. (b) Venn diagram illustrating the overlap of detected proteins between native and dECM groups. (c) Gene Ontology (GO) analysis of Cellular Component categories. (d) GO analysis of Molecular Function categories.

**Supplementary Figure S3.**
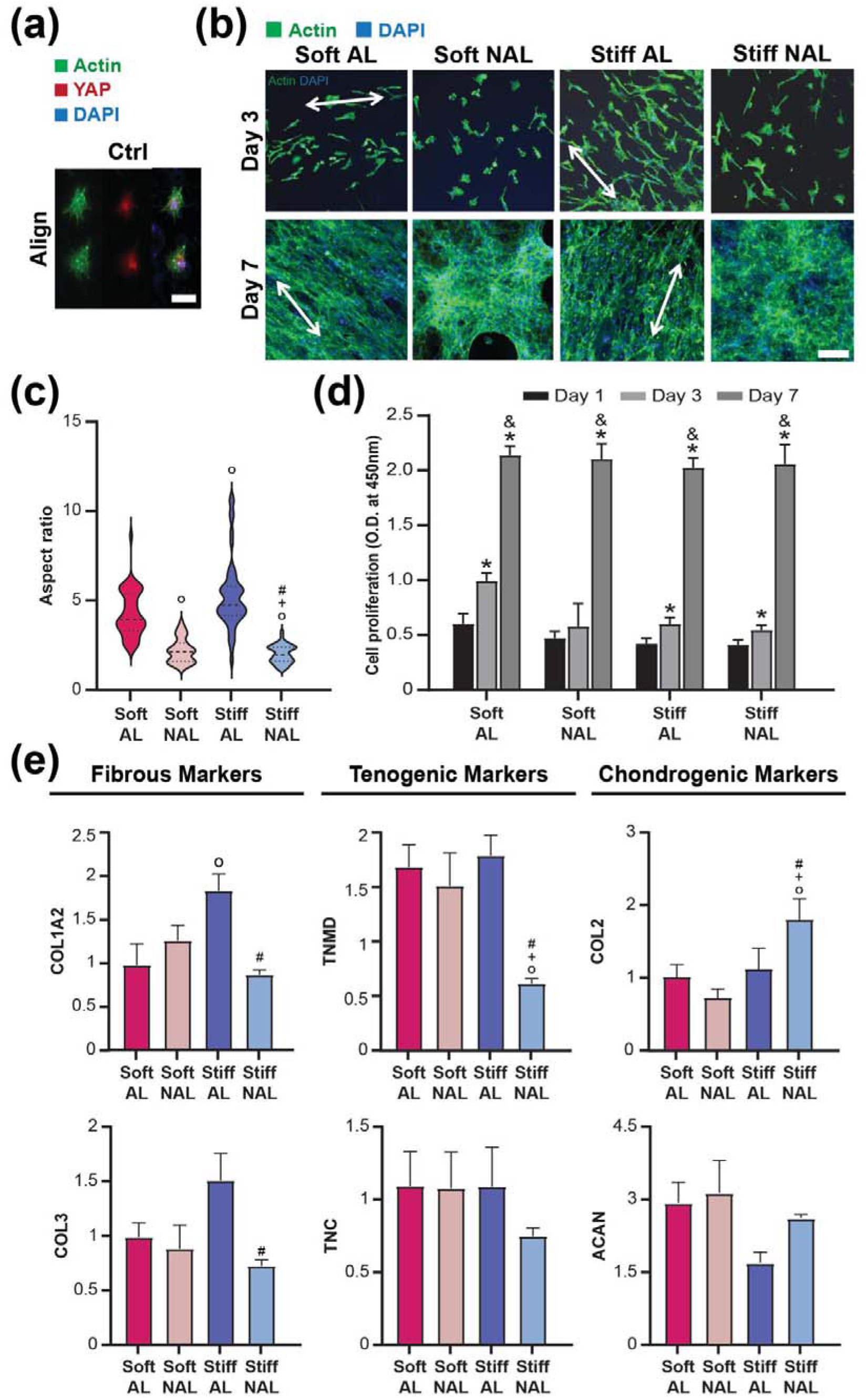
(a) Representative images of bMSCs cultured on aligned 100 % modified MeHA-only scaffolds (Ctrl) for 3 days (Green: F-actin, Red: YAP, Blue: DAPI; Scale bar: 100 µm). (b) Representative bTC images cultured for 3 and 7 days (Allows: fiber direction; Green: F-actin, Blue: DAPI; Scale bar: 600 µm). (c) Quantification of bTC aspect ratios (○: p < 0.0001 vs. Soft AL, +: p < 0.0001 vs. Soft NAL, #: p < 0.0001 vs. Stiff AL, n = 50). (d) bTC proliferation determined by CCK8 assay (*: p < 0.005 vs. Day 1, &: p < 0.005 vs. Day 3, n = 5/group). (e) bTC gene expression at Day 7 (○: p < 0.05 vs. Soft AL, +: p < 0.05 vs. Soft NAL, #: p < 0.05 vs. Stiff AL, n = 5).

**Supplementary Figure S4.**
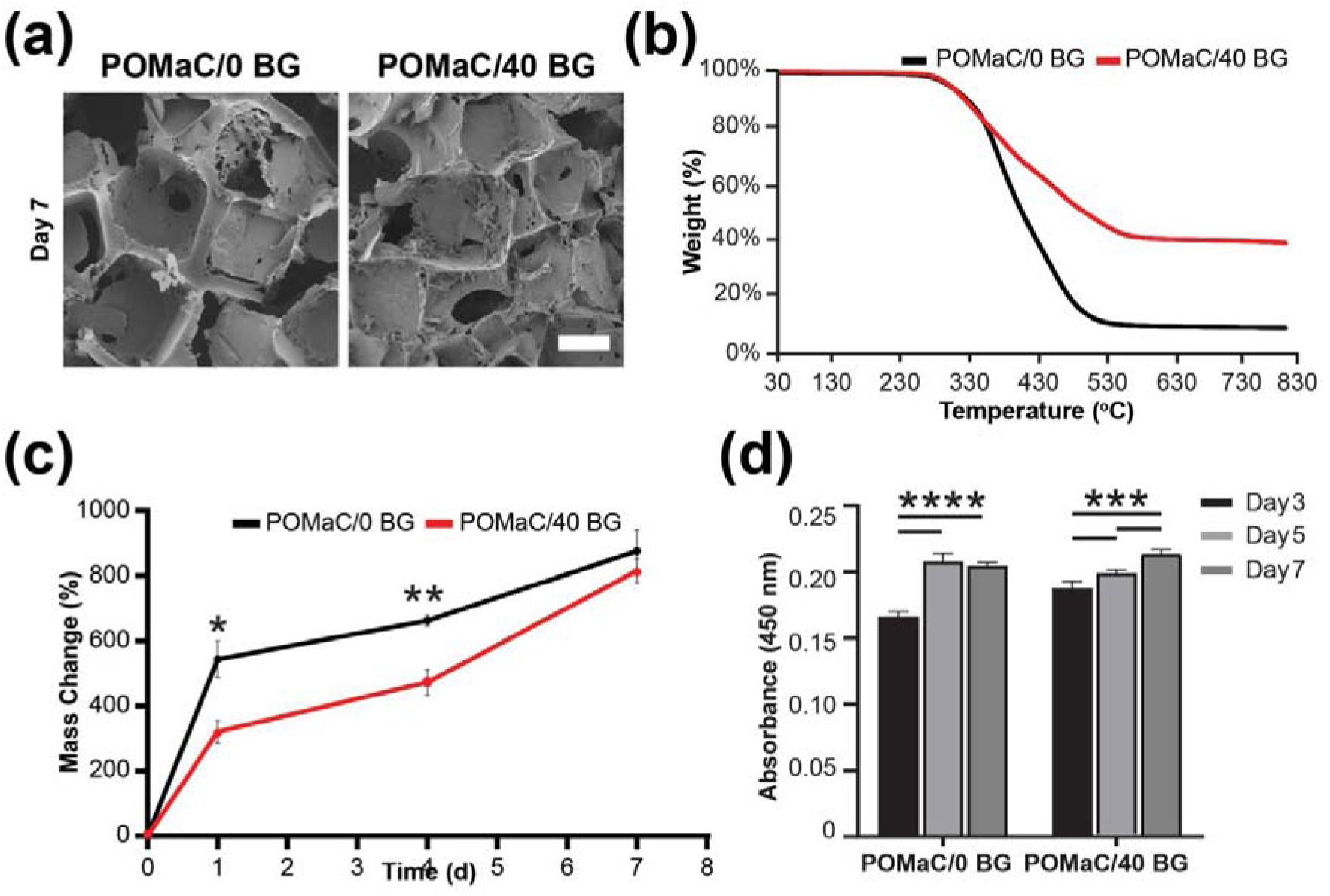
(a) SEM images of POMaC/0 BG and POMaC/40 BG scaffolds on Day 7, showing surface morphology and mineral deposition (scale bar = 200 µm). (b) Thermogravimetric analysis (TGA) of POMaC/0 BG and POMaC/40 BG scaffolds conducted under nitrogen atmosphere from 30°C to 800°C, showing thermal stability and inorganic content. (c) Swelling behavior of scaffolds with significantly higher swelling observed in POMaC/40 BG (*: p < 0.05 at Day 1, **: p < 0.001, n = 4/group). (d) bMSC proliferation assessed via CCK-8 assay, showing enhanced cell growth on POMaC/40 BG scaffolds (p < 0.001, ***p < 0.00001, n = 5/group).

**Supplementary Figure S5.**
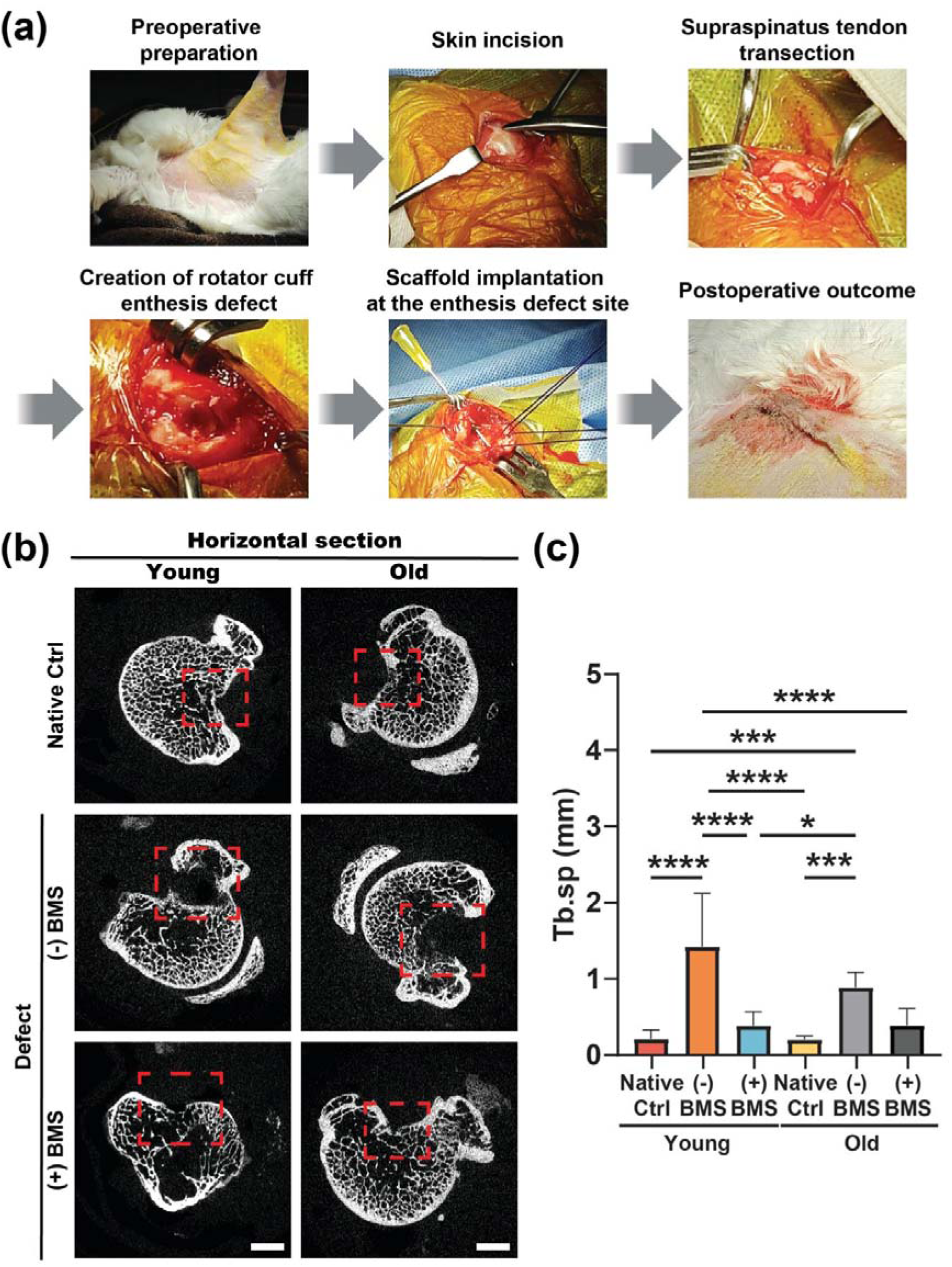
(a) Surgical procedure for scaffold implantation at the rotator cuff enthesis. (b) Horizontally sectioned micro-CT imaging at 4 weeks post-operation, comparing Native Ctrl (native control), [(-) BMS] (defect), and [(+) BMS] (treated) groups, with highlighted regions of interest. (c) Quantitative analysis of the trabecular separation (Tb.Sp) in the regions of interest. Significant differences were observed among groups (n = 4–7; *: p < 0.005, ***: p < 0.0001, ****: p < 0.00001).

**Table S1.**
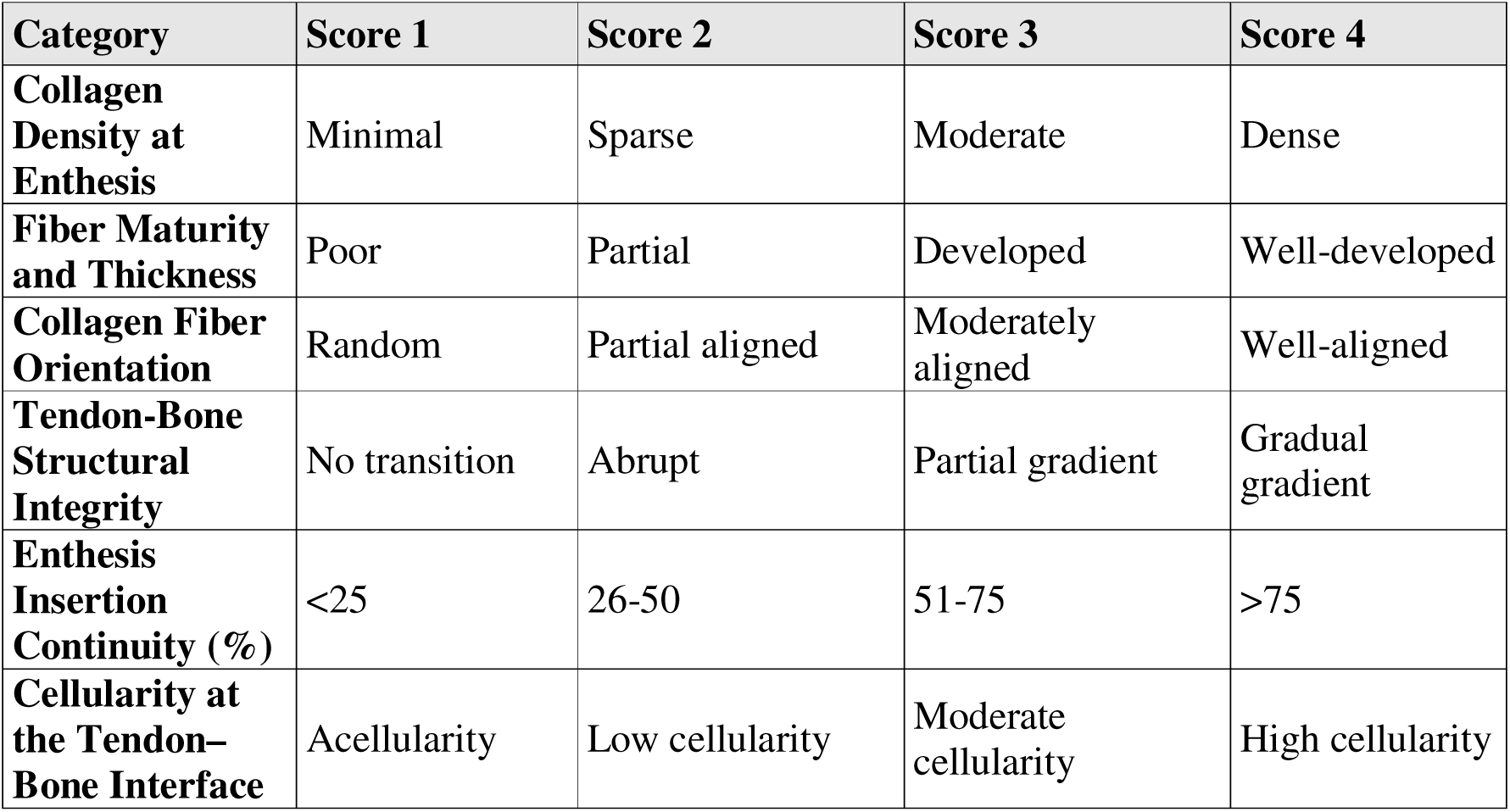
Modified histological scoring system for assessing rotator cuff enthesis regeneration.

**Table S2.**
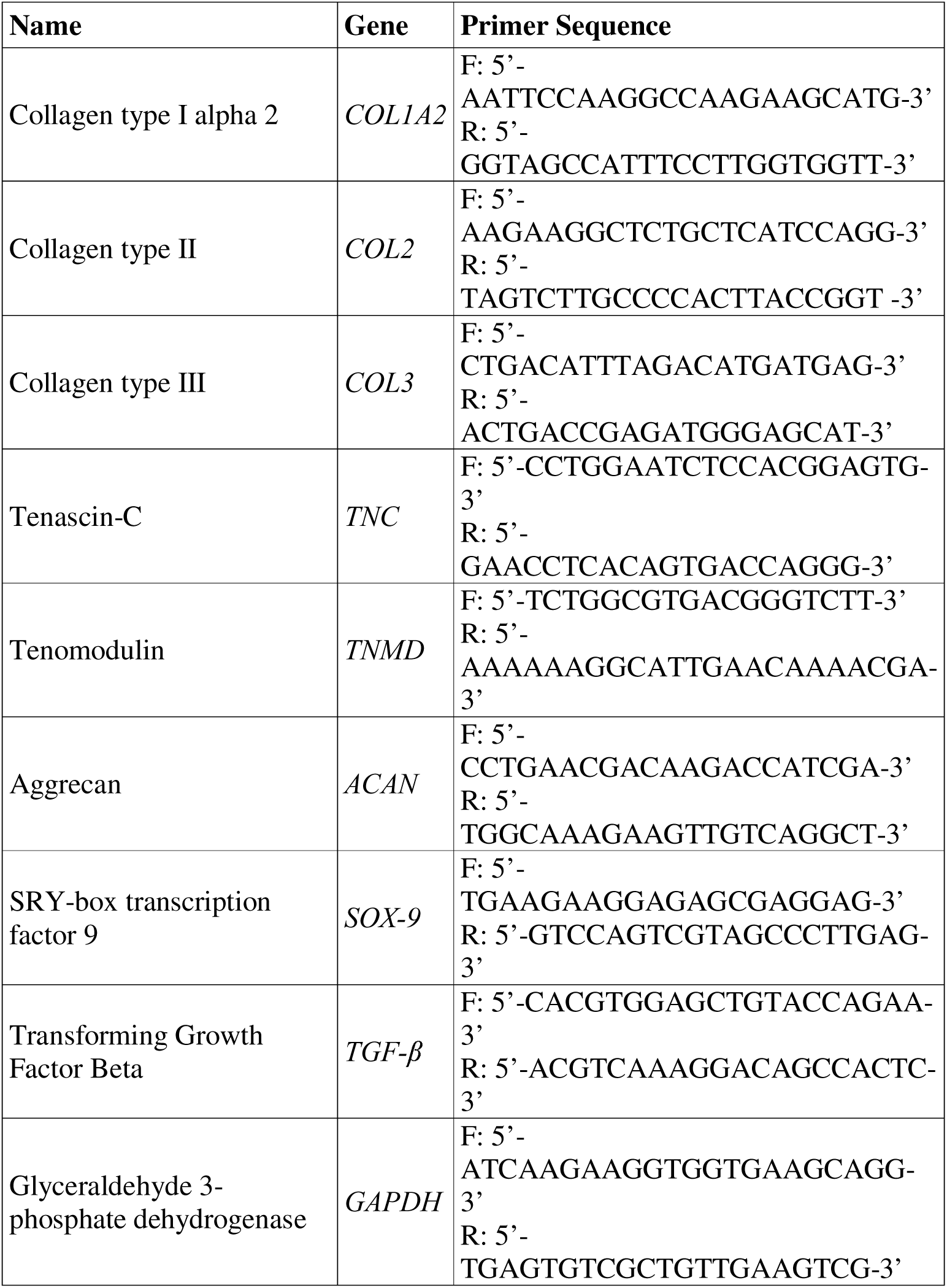
Bovine primers used for RT-qPCR.

