## Supplemental Information for "Engineering a Biomimetic Multiphasic Suture Anchor System for Enhanced Rotator Cuff Enthesis Regeneration"

### Supplementary Information

### Supplementary Materials and Methods

***Chemicals*:** SDS, 4,6-diamidino-2-phenylindole (DAPI), Acetyl-H3K9 (H3K9ac) primary antibody, and Alexa Fluor 488 goat anti-rabbit secondary antibody were purchased from Invitrogen. phosphate-buffered saline (PBS), Dulbecco's minimal essential medium (DMEM) were purchased from Gibco. EDTA was purchased from Thomax Scientific. 60 kDa Sodium hyaluronate was purchased from Lifecore. Methacrylic anhydride, 5M NaOH, Irgacure 2959, 1,8-octanediol, citric acid, maleic anhydride, 1,4-dioxane, 5-(4,6-dichlorotriazinyl) aminofluorescein (DTAF), bovine serum albumin (BSA), and neutral buffered formalin (NBF) were purchased from Sigma Aldrich. 900 kDa poly(ethylene oxide) (PEO) was purchased from Acros Organics. Bioactive glass (BG) was purchased from Mo-Sci Corporation. Sodium chloride, TRIzol, and choloroform were purchased from Fisher Scientific. Fetal bovine serum (FBS) was purchased from R&D systems. Penicillin Streptomycin solution was purchased from Corning. Anti-YAP mouse monoclonal IgG2a antibody was purchased from Santa Cruz Biotechnology. Alexa Fluor-546 goat anti-mouse IgG antibody and Alexa Fluor-488 phalloidin were purchased from Life Technologies Corporation. Cell Counting Kit-8 reagent was purchased from Dojindo. PhotoScript II First-Strand cDNA Synthesis Kit was purchased from New England BioLabs. Fast SYBR Green Master Mix was purchased from Applied biosystems. Graded ethanol was purchased from Decon Labs. Paraffin was purchased from Azer Scientific.

**Supplementary Figures and Legends**

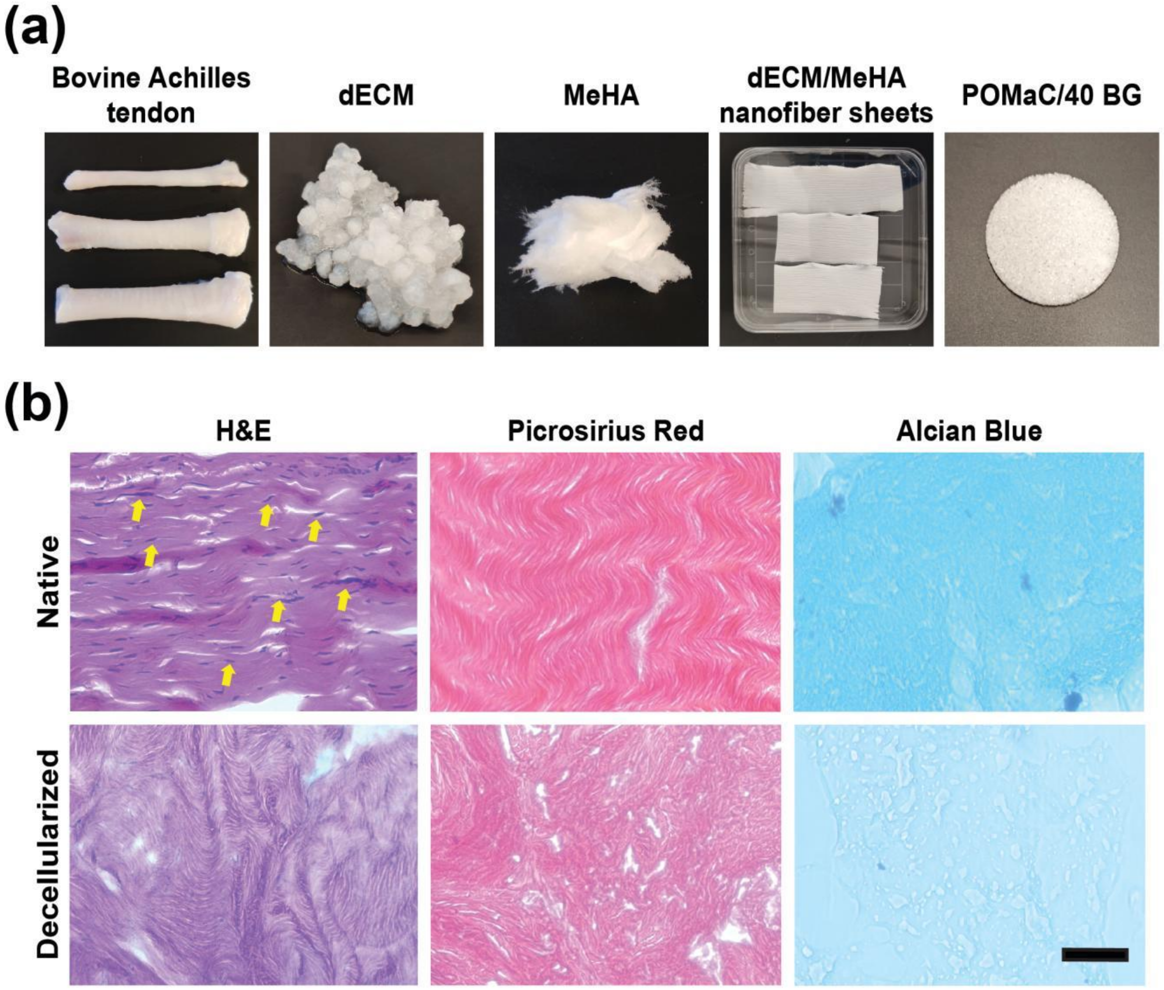


**Supplementary Figure S1**. (a) Representative images of bovine Achilles tendon, decellularized ECM (dECM), stiff MeHA (modification rate = 100%), electrospun dECM/MeHA nanofiber scaffolds (top: stiff aligned (AL); middle: soft non-aligned (NAL); bottom: soft non-aligned (NAL)), and porous POMaC/40% BG scaffold. (b) Representative H&E, Picrosirius Red, and Alcian Blue staining of native and decellularized ECM tissues (Arrows indicate nuclei, scale bar = 50 µm).

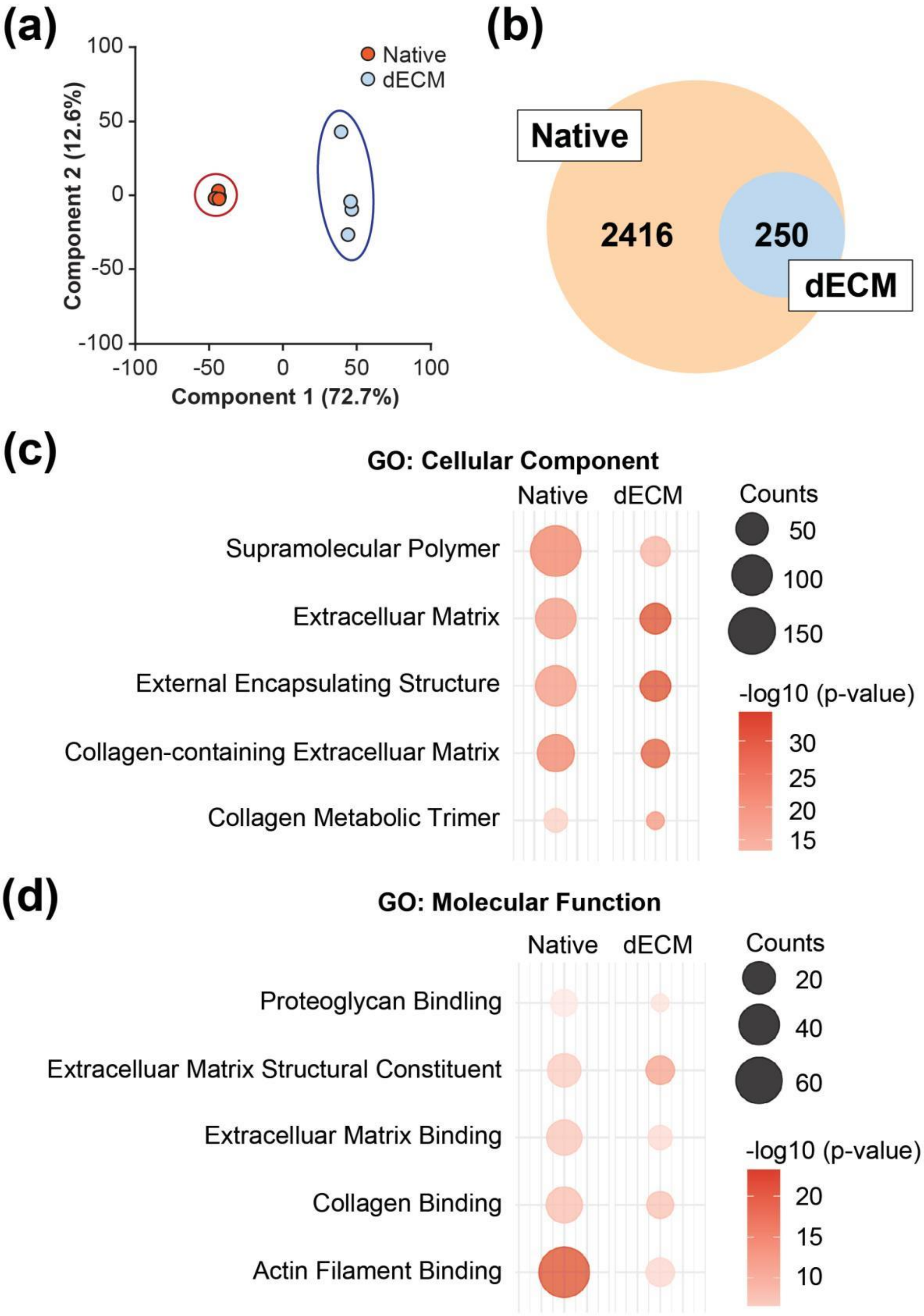


**Supplementary Figure S2**. (a) Principal component analysis (PCA) showing distinct clustering of native and dECM samples. (b) Venn diagram illustrating the overlap of detected proteins between native and dECM groups. (c) Gene Ontology (GO) analysis of Cellular Component categories. (d) GO analysis of Molecular Function categories.

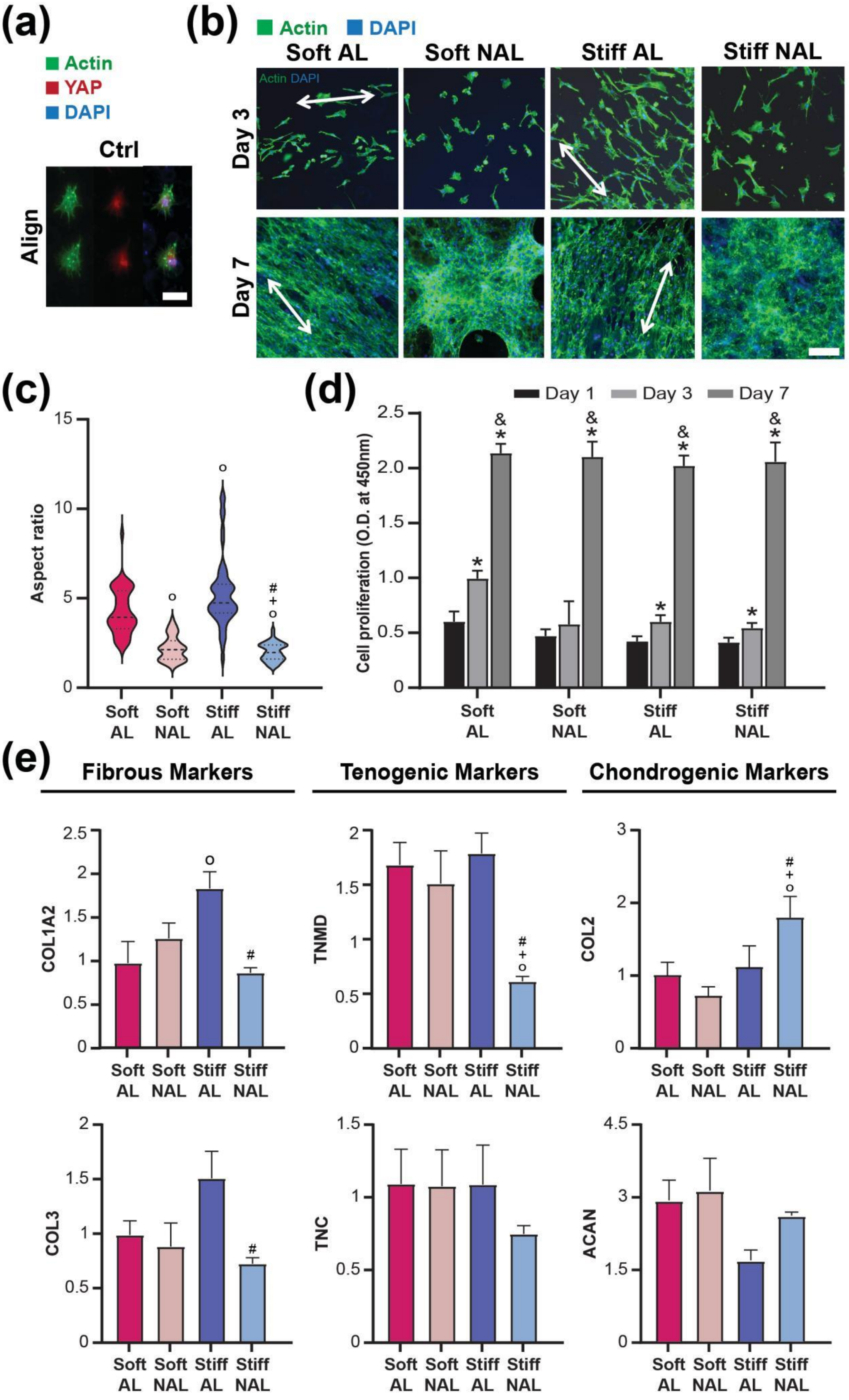

(a)
Actin
YAP
DAPI
Ctrl
Align
(b)
Actin
DAPI
Soft AL
Soft NAL
Stiff AL
Stiff NAL
Day 3
Day 7
(c)
Aspect ratio
Soft AL
Soft NAL
Stiff AL
Stiff NAL
(d)
Day 1
Day 3
Day 7
Cell proliferation (O.D. at 450nm)
Soft AL
Soft NAL
Stiff AL
Stiff NAL
(e)
Fibrous Markers
Tenogenic Markers
Chondrogenic Markers
COL1A2
TNMD
COL2
COL3
TNC
ACAN

**Supplementary Figure S3.** (a) Representative images of bMSCs cultured on aligned 100 % modified MeHA-only scaffolds (Ctrl) for 3 days (Green: F-actin, Red: YAP, Blue: DAPI; Scale bar: 100 μm). (b) Representative bTC images cultured for 3 and 7 days (Allows: fiber direction; Green: F-actin, Blue: DAPI; Scale bar: 600 μm). (c) Quantification of bTC aspect ratios (○: $p < 0.0001$ vs. Soft AL, +: $p < 0.0001$ vs. Soft NAL, #: $p < 0.0001$ vs. Stiff AL, n = 50). (d) bTC proliferation determined by CCK8 assay (*: $p < 0.005$ vs. Day 1, &: $p < 0.005$ vs. Day 3, n = 5/group). (e) bTC gene expression at Day 7 (○: $p < 0.05$ vs. Soft AL, +: $p < 0.05$ vs. Soft NAL, #: $p < 0.05$ vs. Stiff AL, n = 5).

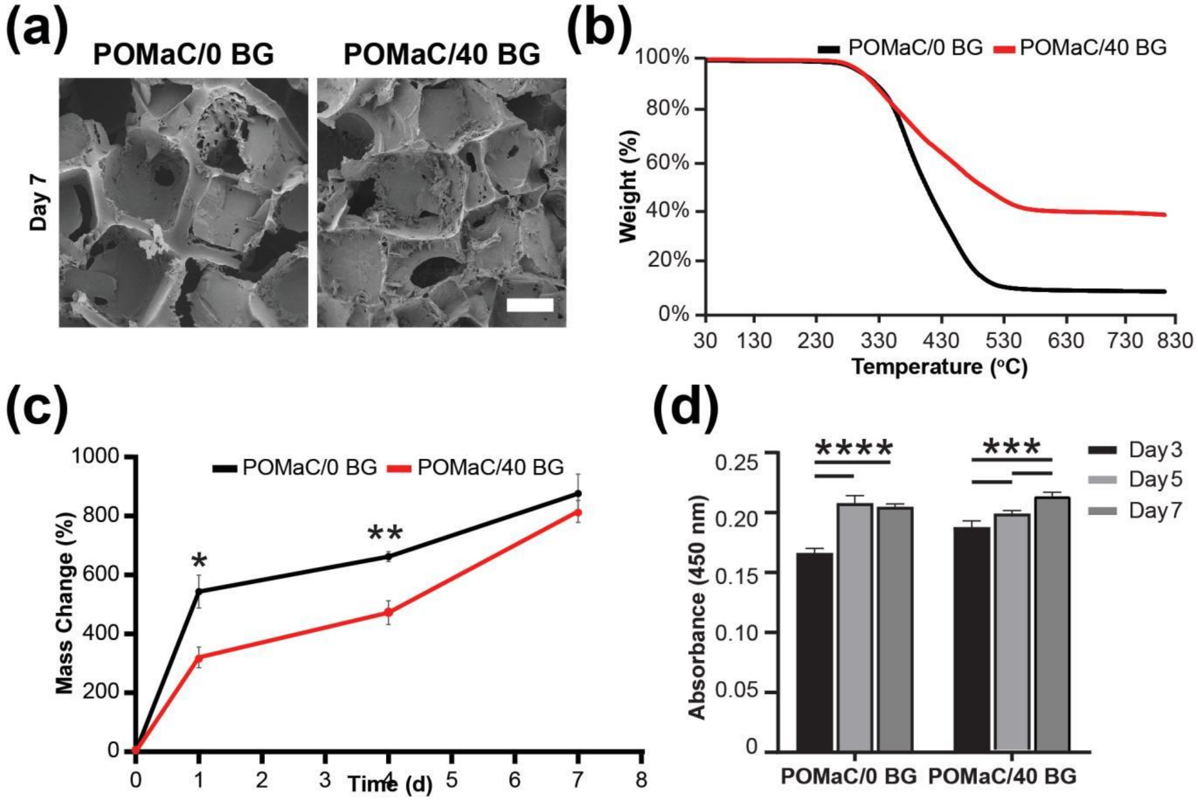


**Supplementary Figure S4**. (a) SEM images of POMaC/0 BG and POMaC/40 BG scaffolds on Day 7, showing surface morphology and mineral deposition (scale bar = 200 µm). (b) Thermogravimetric analysis (TGA) of POMaC/0 BG and POMaC/40 BG scaffolds conducted under nitrogen atmosphere from 30°C to 800°C, showing thermal stability and inorganic content. (c) Swelling behavior of scaffolds with significantly higher swelling observed in POMaC/40 BG (*: $p < 0.05$ at Day 1, **: $p < 0.001$, n = 4/group). (d) bMSC proliferation assessed via CCK-8 assay, showing enhanced cell growth on POMaC/40 BG scaffolds ($p < 0.001$, ***$p < 0.00001$, n = 5/group).

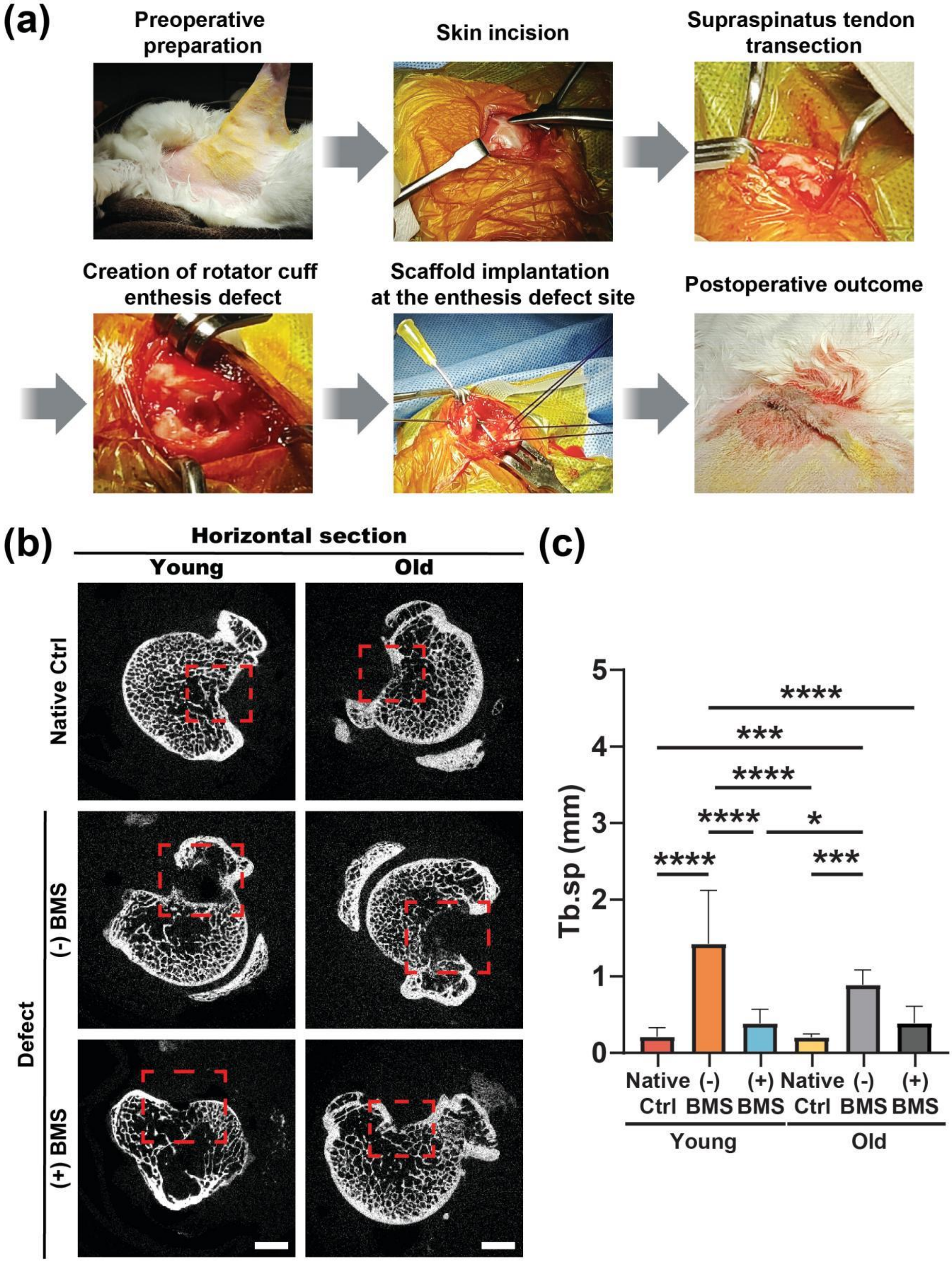


**Supplementary Figure S5**. (a) Surgical procedure for scaffold implantation at the rotator cuff enthesis. (b) Horizontally sectioned micro-CT imaging at 4 weeks post-operation, comparing Native Ctrl (native control), [(-) BMS] (defect), and [(+) BMS] (treated) groups, with highlighted regions of interest. (c) Quantitative analysis of the trabecular separation (Tb.Sp) in the regions of interest. Significant differences were observed among groups ($n = 4–7$; *: $p < 0.005$, ***: $p < 0.0001$, ****: $p < 0.00001$).

**Table S1.** Modified histological scoring system for assessing rotator cuff enthesis regeneration.

| Category | Score 1 | Score 2 | Score 3 | Score 4 |
|---|---|---|---|---|
| **Collagen Density at Enthesis** | Minimal | Sparse | Moderate | Dense |
| **Fiber Maturity and Thickness** | Poor | Partial | Developed | Well-developed |
| **Collagen Fiber Orientation** | Random | Partial aligned | Moderately aligned | Well-aligned |
| **Tendon-Bone Structural Integrity** | No transition | Abrupt | Partial gradient | Gradual gradient |
| **Enthesis Insertion Continuity (%)** | <25 | 26-50 | 51-75 | >75 |
| **Cellularity at the Tendon–Bone Interface** | Acellularity | Low cellularity | Moderate cellularity | High cellularity |

**Table S2.** Bovine primers used for RT-qPCR.

| **Name** | **Gene** | **Primer Sequence** |
|---|---|---|
| Collagen type I alpha 2 | *COL1A2* | F: 5'-AATTCCAAGGCCAAGAAGCATG-3'<br>R: 5'-GGTAGCCATTTCCTTGGTGGTT-3' |
| Collagen type II | *COL2* | F: 5'-AAGAAGGCTCTGCTCATCCAGG-3'<br>R: 5'-TAGTCTTGCCCCACTTACCGGT -3' |
| Collagen type III | *COL3* | F: 5'-CTGACATTTAGACATGATGAG-3'<br>R: 5'-ACTGACCGAGATGGGAGCAT-3' |
| Tenascin-C | *TNC* | F: 5'-CCTGGAATCTCCACGGAGTG-3'<br>R: 5'-GAACCTCACAGTGACCAGGG-3' |
| Tenomodulin | *TNMD* | F: 5'-TCTGGCGTGACGGGTCTT-3'<br>R: 5'-AAAAAAGGCATTGAACAAAACGA-3' |
| Aggrecan | *ACAN* | F: 5'-CCTGAACGACAAGACCATCGA-3'<br>R: 5'-TGGCAAAGAAGTTGTCAGGCT-3' |
| SRY-box transcription factor 9 | *SOX-9* | F: 5'-TGAAGAAGGAGAGCGAGGAG-3'<br>R: 5'-GTCCAGTCGTAGCCCTTGAG-3' |
| Transforming Growth Factor Beta | *TGF-β* | F: 5'-CACGTGGAGCTGTACCAGAA-3'<br>R: 5'-ACGTCAAAGGACAGCCACTC-3' |
| Glyceraldehyde 3-phosphate dehydrogenase | *GAPDH* | F: 5'-ATCAAGAAGGTGGTGAAGCAGG-3'<br>R: 5'-TGAGTGTCGCTGTTGAAGTCG-3' |
